# Orexin signaling in GABAergic lateral habenula neurons modulates aggressive behavior

**DOI:** 10.1101/811265

**Authors:** Meghan E. Flanigan, Hossein Aleyasin, Long Li, C. Joseph Burnett, Kenny L. Chan, Katherine B. LeClair, Elizabeth K. Lucas, Bridget Matikainen-Ankney, William Janssen, Aki Takahashi, Caroline Menard, Madeline L. Pfau, Sam A. Golden, Sylvain Bouchard, Erin S. Calipari, Eric J. Nestler, Ralph J. DiLeone, Akihiro Yamanaka, George W. Huntley, Roger L. Clem, Scott J. Russo

**Affiliations:** Department of Neuroscience, Icahn School of Medicine at Mount Sinai, New York, NY, 10029, USA; Friedman Brain Institute, Icahn School of Medicine at Mount Sinai, New York, NY, 10029, USA; Laboratory of Behavioral Neuroendocrinology, University of Tsukuba, Tsukuba, Ibaraki 305-8577, Japan; Department of Molecular Biomedical Sciences, North Carolina State University, Raleigh, NC, 27695 USA; Department of Psychiatry and Neuroscience, Faculty of Medicine and CERVO Brain Research Center, Université Laval, Ville de Québec, QC G1V-0A6, Canada; Department of Biological Structure, University of Washington, Seattle, Washington, 98185 USA; Department of Psychiatry, Yale University, New Haven, CT, 06519, USA; Department of Neuroscience II, Research Institute of Environmental Medicine, Nagoya University, Nagoya, 464-8601, Japan; Department of Pharmacology, Vanderbilt University School of Medicine, Nashville, TN, 37232, USA

**Author notes:** Corresponding author: Scott J. Russo 1425 Madison Avenue, Room 10-20a New York, NY, 10029, USA.

## Abstract

Heightened aggression is characteristic of multiple neuropsychiatric disorders and can have a wide variety of negative effects on patients, their families, and the public. Recent studies in humans and animals have implicated brain reward circuits in aggression and suggest that, in subsets of aggressive individuals, repeated domination of subordinate social targets is reinforcing. Here, we show that orexin neurons originating from the lateral hypothalamus activate a small population of GABAergic interneurons in the lateral habenula (LHb) via orexin receptor 2 (OxR2) to promote aggression and conditioned place preference (CPP) for aggression-paired contexts. Our study suggests that the orexin system is a potential target for the development of novel therapies aimed at reducing aggressive behaviors and provides the first functional evidence of a local inhibitory circuit within the LHb.

## Main Text

Individuals suffering from a variety of psychiatric syndromes display increased risk for pathological aggressive behaviors^1^. Some of these syndromes include autism spectrum disorders^2^, ADHD^3^, personality disorders^4^, and mood disorders^5^. It has been hypothesized that brain reward systems controlling the valence of social interactions are dysregulated in these individuals, leading to heightened aggression that is reinforcing^6–8^. Recent studies in animals find that subsets of highly aggressive mice will lever press for access to subordinate intruders^9–12^ and form conditioned-place preferences (CPP) for contexts that are associated with access to subordinate intruders^13, 14^. This suggests that mice find the domination of subordinate social targets to be rewarding and display high motivation for repeated opportunities to fight.

Given the strong motivational component to aggressive behavior, there has been increasing interest in the role that reward circuitry plays in controlling aggression. One region in particular, the lateral habenula (LHb), has been newly identified as a potential modulator of the valence of aggressive social interactions^15^. The LHb is a critical node within the reward circuitry of humans and animals that, when activated, promotes negative emotional states predominantly through indirect inhibition of midbrain dopamine neurons^16, 17^. LHb function is disrupted in a variety of neuropsychiatric disorders associated with aggression, including mood disorders^18, 19^. In zebrafish and mice, functional manipulation of LHb neurons alters aggression, its rewarding properties, and the likelihood of “winning” a fight. Specifically, optogenetic inhibition of mouse LHb neurons during the test phase of the aggression CPP task increases the time spent in the aggression-paired context, while optogenetic activation of these neurons reduces it^14^. In addition, direct optogenetic activation of a zebrafish homologue of the mammalian LHb results in increased probability of losing a fight^20^. While these results highlight the importance of the LHb in aggression, the manner in which aggressive social information is integrated by the complex microcircuitry of the LHb is largely unknown.

Previous studies have described the LHb as consisting almost exclusively of glutamate projection neurons expressing vesicular glutamate transporter 2 (vGlut2)^21^. However, there is recent histological and single-cell sequencing evidence that the LHb contains a small population of cells that express glutamate decarboxylase 2 (GAD2)^22–25^, which is generally a marker of GABAergic inhibitory neurons. Despite these recent studies, very little is known about GAD2 neurons in the LHb, including which inputs target them or whether they are capable of providing local inhibition, long-range inhibition, or both. Recent studies have reported that they are enriched in receptors for a number of hypothalamic-derived neuropeptides and hormones, including orexin receptor 2 (OxR2)^23, 25^. As orexin neuron cell bodies are located solely in the lateral hypothalamus (LH)^26^, this suggests that LHb GAD2 neurons are modulated by projections from lateral hypothalamic orexin neurons. Though not previously implicated in aggression, orexin has been strongly implicated in arousal ^27^, social behavior ^28^, and a wide array of motivated behaviors^29–33^. Here, we set out to characterize the cell type-specific dynamics of LHb neurons during aggression and determine whether orexin release onto GAD2 LHb neurons affects aggressive behavior. First, we found that while LHb vGlut2 neurons reduce their activity during aggression, GAD2 LHb neurons increase their activity. Direct optogenetic manipulation of GAD2 LHb neurons, which exert local inhibitory control over LHb neurons, alters aggression and aggression CPP. Further, we demonstrate that direct modulation of orexin inputs from the LH to the LHb promotes aggression and aggression CPP through an OxR2-dependent mechanism. Our findings indicate that cell-type specific engagement of orexin signaling in the LHb is important for modulating aggression and extend orexin’s known role in motivation to the social realm.

## Results

### LHb neural responses to social targets

To first investigate patterns of total LHb activity associated with repeated aggressive social encounters, we injected non-conditional AAV-GCaMP6 into the LHb, implanted a fiber above the cells, and measured fluorescent calcium transients in highly aggressive (AGG) and non-aggressive (NON) CD-1 wild-type mice during the resident intruder (RI) and aggression CPP tests using fiber photometry (Fig. 1a-b). In the RI test, male CD-1 outbred mice are exposed to submissive male C57BL/6J intruders for 5 minutes in their home cage and allowed to freely interact. CD-1 mice display individual variation in aggressive behaviors in this task, with some animals consistently fighting with low attack latencies (termed aggressors, AGGs) and some animals never engaging in any aggressive behaviors (termed non-aggressors, NONs). Importantly, because CD-1 resident mice are vastly larger and more dominant than C57BL6/J intruder mice, AGGs engaging in fighting during RI consistently subordinate intruder mice and are never themselves attacked by the intruder. In aggression CPP, animals are conditioned in two 10 minute sessions per day for 3 days to associate distinct contexts with the presence or absence of subordinate intruders. On test day, experimental mice are given free access to both contexts (in the absence of intruders) and the time spent in each context is measured to determine the valence of the social encounter. NONs form an aversion for the intruder-paired context, whereas AGGs form a preference for it, suggesting that AGGs find these social encounters rewarding and will seek out opportunities for aggression^14^.

**Figure 1:**
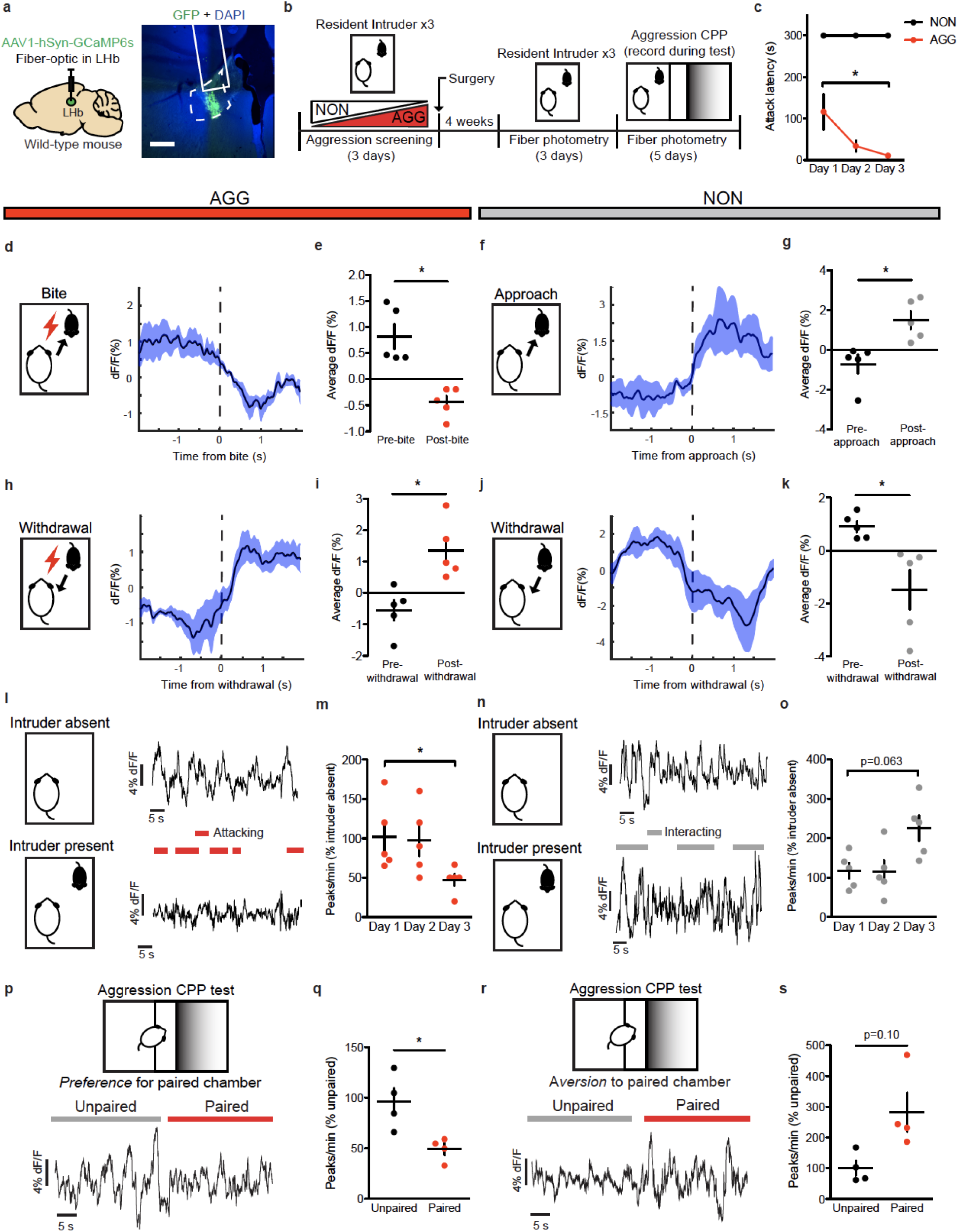
Aggressive behaviors are associated with decreased LHb activity. **a,** Surgical manipulations and representative viral infection for LHb photometry experiments, scale bar = 400 μm. **b,** Experimental timeline for LHb photometry experiments. **c,** AGGs displayed reduced attack latency on day 3 compared to day 1 of RI (one-way repeated measures ANOVA, n=5 AGGs, F(2,17)=7.248, main effect of day p=0.0113, Bonferroni posthoc d1 vs d3, p<0.05). **d,** Peri-event plot of AGG LHb activity before and after a bite on day 3 of RI. For all peri-event plots, black line denotes the mean signals for all animals, while blue shaded region denotes the SEM. **e,** AGGs displayed a reduction in average LHb activity following a bite (paired t-test, n=5 mice, 3-5 bites per mouse, t(4)=4.593, p=0.0101). **f,** Peri-event plot of NON LHb activity before and after an intruder approach on day 3 of RI. **g,** NONs displayed an increase in average LHb activity following an approach (paired t-test, n=5 mice, 3-5 approaches per mouse, t(4)=3.450, p=0.0261. **h,** Peri-event plot of AGG LHb activity before and after a withdrawal from an aggressive bout. **i,** AGGs displayed an increase in average LHb activity following a withdrawal (paired t-test, n=5 mice, 3-5 withdrawals per mouse, t(4)=3.911, p=0.0174. **j,** Peri-event plot of NON LHb activity before and after a withdrawal from a non-aggressive social interaction. **k,** NONs displayed a decrease in LHb activity after a withdrawal from a non-aggressive social interaction (paired t-test, n=5 mice, 3-5 withdrawals per mouse, t(4)=2.838, p=0.0470). **l,** Representative traces of AGG LHb activity in the absence and presence of an intruder mouse on day 3 of RI. **m,** AGGs displayed reduced LHb activity across three days of RI (one-way repeated measures ANOVA, n=5 mice, F(2,14)=6.294, p=0.0228, Bonferroni posthoc test, day 1 vs day 3 p<0.05). **n,** Representative traces of NON LHb activity in the absence and presence of an intruder mouse on day 3 of RI. **o,** NONs displayed a trend towards increased LHb activity across 3 days of RI (one-way repeated measures ANOVA, n=5 mice, F(2,14)=3.985, p=0.0630). **p,** Representative trace of AGG LHb activity during the aggression CPP test. **q,** AGGs displayed reduced LHb activity in the paired context compared to the unpaired context during the aggression CPP test (paired t-test, n=4 mice, t(3)=4.080, p=0.0266). **r,** Representative trace of NON LHb activity during the aggression CPP test. **s,** NONs displayed no differences in LHb activity in the paired context compared to the unpaired context during the aggression CPP test (paired t-test, n=4 mice, t(3)=2.352, p=0.1001). *p<0.05. All data are expressed as mean ± SEM.

Given that optogenetic inhibition of the LHb has been shown to promote aggression and aggression CPP^14^, we hypothesized that aggressive behaviors would be associated with reductions in LHb activity. Consistent with our hypothesis, AGGs displayed reductions in LHb activity upon biting an intruder (Fig. 1d-e). Conversely, NONs displayed increases in LHb activity upon intruder approaches (Fig. 1f-g). AGGs and NONs also displayed opposing LHb responses to withdrawals from social bouts, with AGGs showing increases in LHb activity upon withdrawals from aggressive social bouts and NONs showing decreases in LHb activity upon withdrawals from non-aggressive social bouts (Fig. 1h-k). Importantly, these LHb responses to social targets during RI in AGGs were observed on both day 3 of RI (Fig. 1) and day 1 of RI (Fig. S1). Over the 3 days of RI, AGGs increased their aggression towards the intruder, attacking with shorter latencies on day 3 than on day 1 (Fig. 1c). This is consistent with previous studies describing a phenomenon called “the winner effect,” whereby prior winning experience increases aggression towards future social targets ^34^. Notably, the observed increase in aggression on day 3 of RI was associated with decreased LHb activity across the RI session compared to day 1 of RI (Fig. 1l-m). However, NONs only trended towards increased LHb activity across the RI session on day 3 compared to day 1 (p=0.08, Fig. 1n-o). To determine whether the LHb is capable of encoding contextual information about the rewarding properties of aggressive social encounters, we next performed fiber photometry in AGGs and NONs during the test phase of aggression CPP. We found that AGGs displayed reduced LHb activity in the paired context of the CPP chamber compared to the unpaired context (Fig. 1p-q). Similarly, NONs showed a trend towards increased LHb activity in the paired context of the CPP chamber compared to the unpaired context (p=0.10, Fig. 1r-s). In addition, CPP scores of AGGs and NONs were negatively correlated with LHb activity in the paired context (Fig. S1). Together, these data indicate that the LHb is involved in encoding the rewarding properties of aggressive social interactions and the contexts associated with them.

Although the LHb consists predominantly of glutamatergic projection neurons expressing vGlut2^21^, GAD2-expressing LHb cells have also been reported^22, 24, 25, 35^. To confirm this, we performed *in-situ* hybridization (ISH) for GAD2 mRNA in the LHb (Fig. S2). We found that GAD2 neurons make up ∼18% of total LHb neurons and are localized primarily to the medial aspect of the LHb. To better understand the role of specific LHb cell types (i.e. GAD2 versus vGlut2) in aggressive behavior, we screened mice in RI and compared the number of Fos-positive GAD2 and vGlut2 nuclei in the LHb of AGGs and NONs (Fig. 2a-b). Following the last RI experience, both total Fos-positive nuclei and vGlut2 Fos-positive nuclei were reduced in AGGs compared to NONs (Fig. 2c,e). Interestingly, GAD2 Fos-positive nuclei were increased in AGGs compared to NONs (Fig. 2c-d), suggesting that these cells may be playing a role in regulating aggressive behavior. To assess the dynamics of GAD2 neurons in real-time, we used cell-type specific fiber photometry to record their activity during RI and aggression CPP (Fig. 3a-b). To do this, we injected AAV-Flex-GCaMP6 into the LHb of GAD2-Cre mice and implanted a fiber above the infected cells. Upon biting an intruder, AGGs displayed robust activation of GAD2 neurons (Fig. 3d-e), whereas GAD2 neuron activity decreased upon withdrawal from an aggressive bout (Fig 3h-i). These responses were observed on both day 1 and day 3 of RI, and increases in overall GAD2 neuron activity coincided with increases in aggressive behavior (Fig. S3 and 3c,l-m). Interestingly, the activity of GAD2 neurons in NONs was only weakly associated with social behavior in RI (Fig. 3f-g, j-k), indicating that NON LHb responses to social targets may not be as strongly driven by alterations in the activity of LHb GAD2 neurons. During the aggression CPP test, AGGs displayed increased LHb GAD2 neuron activity in the aggression-paired context compared to the unpaired context (Fig. 3p-q). Though we did not observe any significant differences in LHb GAD2 neuron activity between the paired and unpaired chambers in NONs (Fig. 3r-s), LHb GAD2 neuron activity was correlated with aggression CPP score in both AGGs and NONs (Fig. S3). Therefore, our findings imply that LHb GAD2 neurons represent a sub-population within the LHb whose activity during aggression and exposure to aggression-associated contexts opposes that of the LHb as a whole.

**Figure 2:**
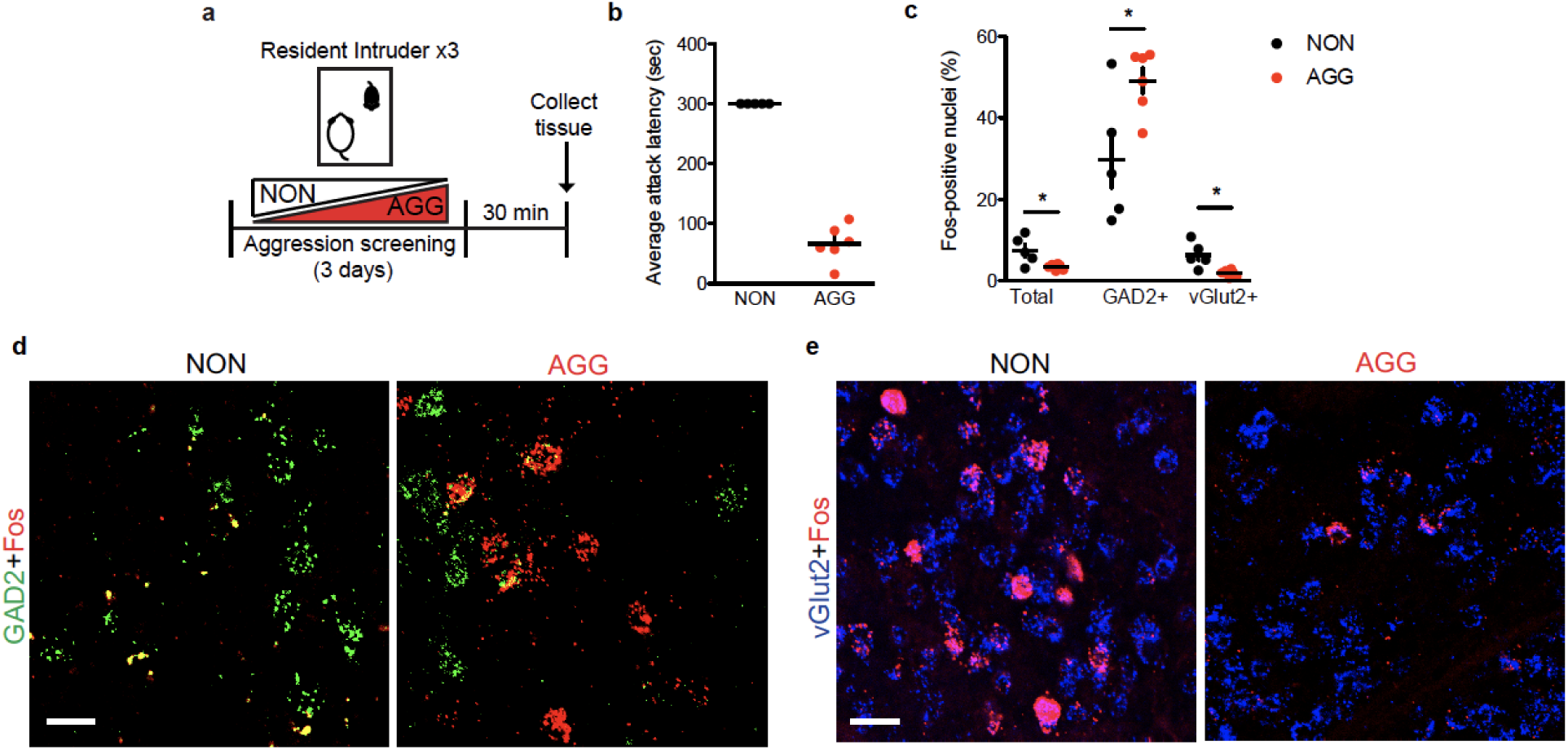
Aggressive behavior is associated with increased expression of Fos mRNA in GAD2 LHb neurons. **a,** Experimental timeline for Fos *in-situ* hybridization (ISH) experiments. **b,** Average attack latency for NONs and AGGs over three days of RI screening. **c,** Fos-positive nuclei were lower in AGGs than in NONs for total and vGlut2-positive LHb neurons and higher in GAD2-positive LHb neurons (student’s t-test, n=5 NONs, n=6 AGGs, 2 slices per mouse, total: t(9)=2.828, p=0.0198, vGlut2: t(9)=3.421, p=0.0176, GAD2: t(9)=2.686, p=0.025). **d,** Representative images of ISH in the LHb for Fos (red) and GAD2 (green) in NON and AGG mice following RI screening, scale bar=35 μm**. e,** Representative images of ISH in the LHb for Fos (red) and vGlut2 (blue) in NON and AGG mice following RI screening, scale bars=35 μm. *p<0.05. All data are expressed as mean ± SEM.

**Figure 3:**
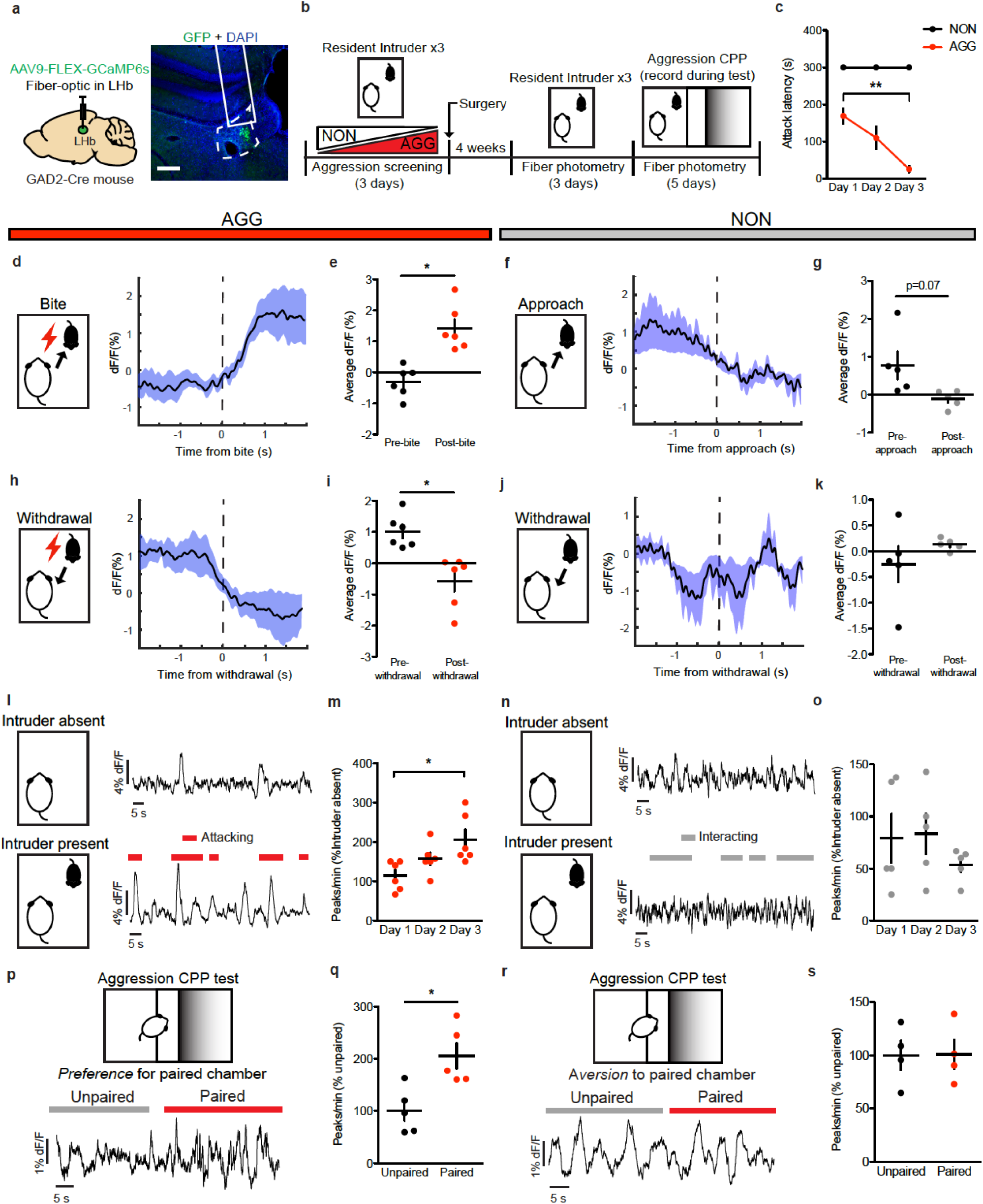
Aggressive behaviors are associated with increased GAD2 LHb neuron activity. **a,** Surgical manipulations and representative viral infection for LHb GAD2 neuron photometry experiments, scale bar=400 μm. **b,** Experimental timeline for LHb GAD2 neuron photometry experiments. **c,** AGGs displayed reduced attack latency on day 3 of RI compared to day 1 of RI (one-way repeated measures ANOVA, n=5 AGGs, F(2,17)=10.78, p=0.0032, main effect of day, Tukey posthoc test, d1 vs d3, p<0.01). **d,** Peri-event plot of AGG LHb GAD2 activity 2s before and after a bite on day 3 of RI. For all peri-event plots, black line denotes the mean signals for all animals, while blue shaded region denotes the SEM. **e,** AGGs displayed an increase in average GAD2 neuron activity following a bite (paired t-test, n=5 mice, 3-5 bites per mouse, t(5)=4.914, p=0.0044). **f,** Peri-event plot of NON LHb GAD2 neuron activity 2s before and after an intruder approach on day 3 of RI. **g,** NONs displayed no change in average LHb activity following an approach (paired t-test, n=5 mice, 3-5 approaches per mouse, t(4)=2.437, p=0.0714. **h,** Peri-event plot of AGG LHb GAD2 neuron activity 2s before and after a withdrawal from an aggressive bout. **i,** AGGs displayed a decrease in average LHb activity following a withdrawal (paired t-test, n=6 mice, 3-5 withdrawals per mouse, t(5)=3.022, p=0.0294. **j,** Peri-event plot of NON LHb GAD2 neuron activity 2s before and after a withdrawal from a non-aggressive social interaction. **k,** NONs displayed no change in LHb GAD2 neuron activity after a withdrawal from a non-aggressive social interaction (paired t-test, n=5 mice, 3-5 withdrawals per mouse, t(4)=1.170, p=0.3068). **l,** Representative traces of AGG LHb GAD2 neuron activity in the absence and presence of an intruder mouse during day 3 of RI. **m,** AGGs displayed increased GAD2 LHb activity across three days of RI (one-way repeated measures ANOVA, n=6 mice, F(2,17)=5.653, p=0.0228, Bonferroni posthoc test day 1 vs. day 3 p<0.05) **n,** Representative traces of NON LHb GAD2 neuron activity in the absence and presence of an intruder mouse during day 3 of RI. **o,** NONs did not display changes in LHb GAD2 neuron activity across three days of RI (one-way repeated measures ANOVA, n=5 mice, F(2,14)=0.8904, p=0.4476). **p,** Representative trace of AGG LHb GAD2 neuron activity during the aggression CPP task. **q,** AGGs displayed increased LHb GAD2 neuron activity in the paired context compared to the unpaired context during the aggression CPP task (paired t-test, t(4)=2.885, p=0.0448). **r,** Representative trace of NON LHb GAD2 neuron activity during the aggression CPP task. **s,** NONs did not display differences in LHb GAD2 neuron activity during the aggression CPP task (paired t-test, t(3)=0.03591, p=0.9736). *p<0.05, **p<0.01. All data are expressed as mean ± SEM.

### Optogenetic manipulation of LHb inhibitory neurons

Though LHb GAD2 neurons have been reported to express GABA as well as the vesicular GABA transporter (vGAT)^25^, it remains unknown whether these neurons promote local inhibition within the LHb, outside of the LHb, or both. To assess whether LHb GAD2 neurons can provide local inhibition, we transduced LHb GAD2 neurons with channel-rhodopsin (ChR2) by injecting AAV-DIO-ChR2 into the LHb of GAD2-Cre mice and applied blue light to brain slices containing the LHb while recording from putative LHb vGlut2 neurons using whole-cell electrophysiology (Fig. 4a). We found that optogenetic stimulation of LHb GAD2 neurons in slice elicited monosynaptic inhibitory currents in non-GAD2 (vGlut2) neurons that were completely blocked with the GABA receptor antagonist picrotocxin (Fig. 4b-c). We did not detect excitatory currents in any neurons as a result of GAD2 neuron optogenetic stimulation, as evidenced from a lack of responses when holding neurons at −70 mV. Moreover, DREADD-mediated activation of LHb GAD2 neurons *in-vivo* reduced the total number of putative non-GAD2 Fos-positive neurons in the LHb (Fig. 4d-g). Together, these data provide the first evidence of a functional inhibitory microcircuit within the LHb and challenge the longstanding notion that the LHb is devoid of local sources of inhibition. To test whether LHb GAD2 neurons project outside of the LHb, we performed anterograde tracing of their projections by injecting AAV-DIO-eYFP into the LHb of GAD2-Cre mice. We did not observe evidence of eYFP-positive axons in primary known LHb target regions like the ventral tegmental area (VTA), the dorsal raphe nucleus (DRN), the median raphe nucleus (MRN), or the rostromedial tegmental nucleus (RMTg) (Fig. S4). These results suggest that LHb GAD2 neurons are capable of inhibiting neurons within the LHb itself and may not directly target neurons outside of the LHb.

**Figure 4:**
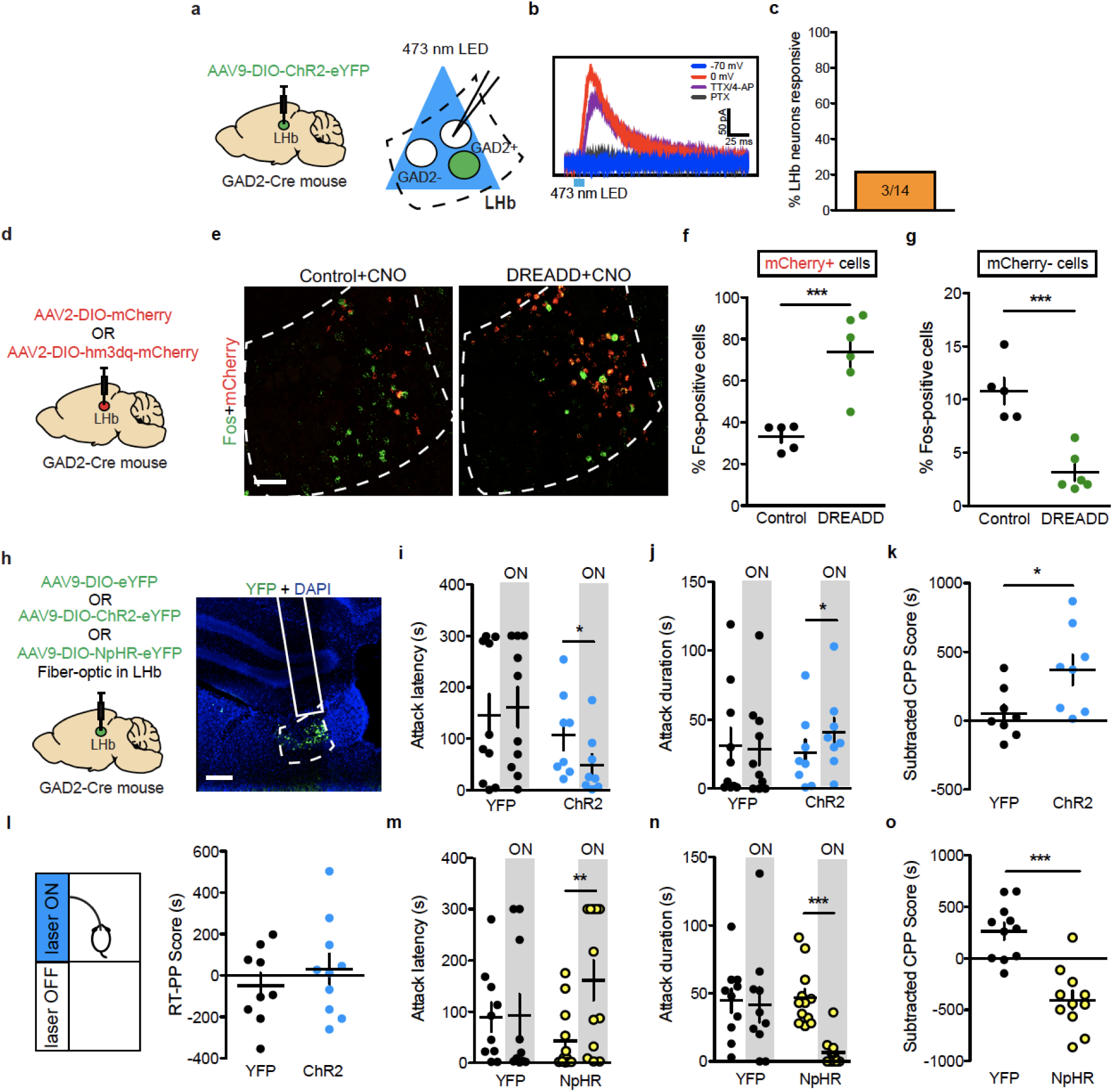
GAD2 LHb neurons are locally inhibitory and regulate aggressive behaviors. **a,** Surgical manipulations and experimental schematic for slice optogenetic stimulation of GAD2 LHb neurons. **b,** Representative trace of GFP-negative neuron in response to optogenetic stimulation of nearby GAD2 neurons when cells are held at 0 mV or −70 mV and in the presence of TTX or the GABA receptor antagonist picrotoxin. **c,** Percent LHb GFP-negative neurons responding to optogenetic stimulation of GAD2 neurons. **d,** Surgical manipulations for GAD2 DREADD experiments. **e,** Representative images of ISH for Fos (green) and mCherry (red) mRNA in control and DREADD mice following treatment with CNO, scale bar= 50μm. **f,** CNO treatment increased the percent of Fos-expressing mCherry-positive neurons in DREADD mice compared to controls (student’s t-test, n=5 mice per group, t(9)=4.905, p=0.0008). **g,** CNO treatment reduced the percent of Fos-expressing mCherry-negative neurons in DREADD mice compared to controls (student’s t-test, n=5 mice per group, t(9)=5.439, p=0.0004). **h,** Surgical manipulations and representative viral infection image for *in-vivo* GAD2 LHb neuron optogenetics experiments, scale bar= 300μm **i,** ChR2-mediated stimulation of GAD2 LHb neurons reduced attack latency in RI (paired t-test, n=8 mice, t(7)=3.724, p=0.0136). **j,** ChR2-mediated stimulation of LHb neurons increased attack duration in RI (paired t-test, n=8 mice, t(7)=2.690, p=0.0311). **k,** ChR2-mediated stimulation of GAD2 LHb neurons increased aggression CPP (student’s t-test, n=8 mice per group, t(14)=2.482, p=0.0264). *p<0.05. **l,** Stimulation of GAD2 LHb neurons did not induce a real-time place preference (student’s t-test, t(17)=0.8271, p=0.4196). **m,** NpHr-mediated inhibition of GAD2 LHb neurons increased attack latency in RI (paired t-test, n=12 mice, t(11)=3.242, p=0.0078). **n,** NpHr-mediated inhibition of GAD2 LHb neurons reduced attack duration in RI (paired t-test, n=12 mice, t(11)=5.504, p=0.002). **o,** NpHr-mediated inhibition of GAD2 LHb neurons reduced aggression CPP (student’s t-test, n=11 mice per group, t(20)=5.517, p<0.0001). *p<0.05, **p<0.01, ***p<0.001. All data are expressed as mean ± SEM.

To determine whether LHb GAD2 neurons play a functional role in aggressive behavior, we performed *in-vivo* optogenetic manipulation of LHb GAD2 neurons during RI and the aggression CPP test (Fig. 4h). To do this, we injected AAV-DIO-ChR2 (channel-rhodopsin) or AAV-DIO-eYFP into the LHb of GAD2-Cre mice and implanted a fiber above the LHb. We found that optogenetic activation of GAD2 LHb neurons with ChR2 in AGG mice (20 Hz, 20 ms pulses, 7mW) reduced the latency to attack and increased the total duration of time spent attacking in RI (Fig. 4i-j). Notably, we were unable to elicit aggression in NONs with optogenetic stimulation of GAD2 neurons (Fig. S5). Optogenetic activation of GAD2 LHb neurons in AGGs during the aggression CPP test also increased the time spent in the intruder-paired context (Fig. 4k). However, stimulation of GAD2 neurons did not elicit a real-time place preference (RTPP) (Fig. 4l) or promote CPP for palatable food (Fig. S6). These data imply that GAD2 neurons potentiate aggression by enhancing its rewarding properties, but do not modulate food reward or general reward-like responding. Consistent with these findings, consumption of palatable food did not alter the activity of LHb GAD2 neurons as determined using fiber photometry and the calcium sensor GCaMP6 (Fig. S6).

To determine if LHb GAD2 neurons are necessary for aggression, we tested whether optogenetic inhibition of them reduces aggression and aggression CPP. To do this, we injected AAV-DIO-NpHR (halo-rhodopsin) or AAV-DIO-eYFP into the LHb of GAD2-Cre mice and implanted a fiber above the LHb. Though this did not completely block aggression, optogenetic inhibition of GAD2 LHb neurons in AGGs (constant light, 8s on/2s off, 7 mW) increased the latency to attack and decreased the total duration of time spent attacking in RI (Fig. 4m-n). Optogenetic inhibition of LHb GAD2 neurons also promoted aversion for the intruder-paired context during aggression CPP (Fig. 4o). Together, these results indicate that LHb GAD2 neurons can alter the valence of aggressive social interactions and subsequently, the intensity of aggression itself.

### Orexin modulation of LHb inhibitory neurons

Though little is currently known about LHb GAD2 neurons, reports suggest they express a number of hypothalamus-derived neuropeptide and hormone receptors, including OxR2^23, 25^. Orexin has not been previously implicated in aggression, but has been implicated in a wide variety of motivated behaviors from drug addiction to social interaction^25–29^. Therefore, we hypothesized that orexin-mediated activation of LHb GAD2 neurons via OxR2 would promote aggressive behavior. First, we characterized the expression of OxR2 in the LHb by performing double fluorescent *in situ* hybridization (ISH) for OxR2 in GAD2 and vGlut2 neurons (Fig. 5a). We detected expression of OxR2 in nearly 100% of LHb GAD2 neurons, while fewer than 10% of LHb vGlut2 neurons were positive for OxR2 (Fig. 5b-c). To visualize orexin-positive axons in the LHb in close proximity to LHb GAD2 neurons, we utilized the high-resolution microscopy technique AiryScan in conjunction with viral-mediated fluorescent labeling (AAV-DIO-GFP in GAD2-Cre mice) of GAD2 cells and immunohistochemistry (IHC) to visualize orexin axons. Following 3D rendering of the images, we observed orexin axons closely apposed to GAD2 cell bodies and processes (Fig. S7). To functionally test the hypothesis that orexin modulates the activity of GAD2 neurons, we performed bath application of orexin in slices containing the LHb. We found that this increased the firing rate of GAD2 neurons, indicating that they are indeed physiologically activated by orexin (Fig. 5d-f).

**Figure 5:**
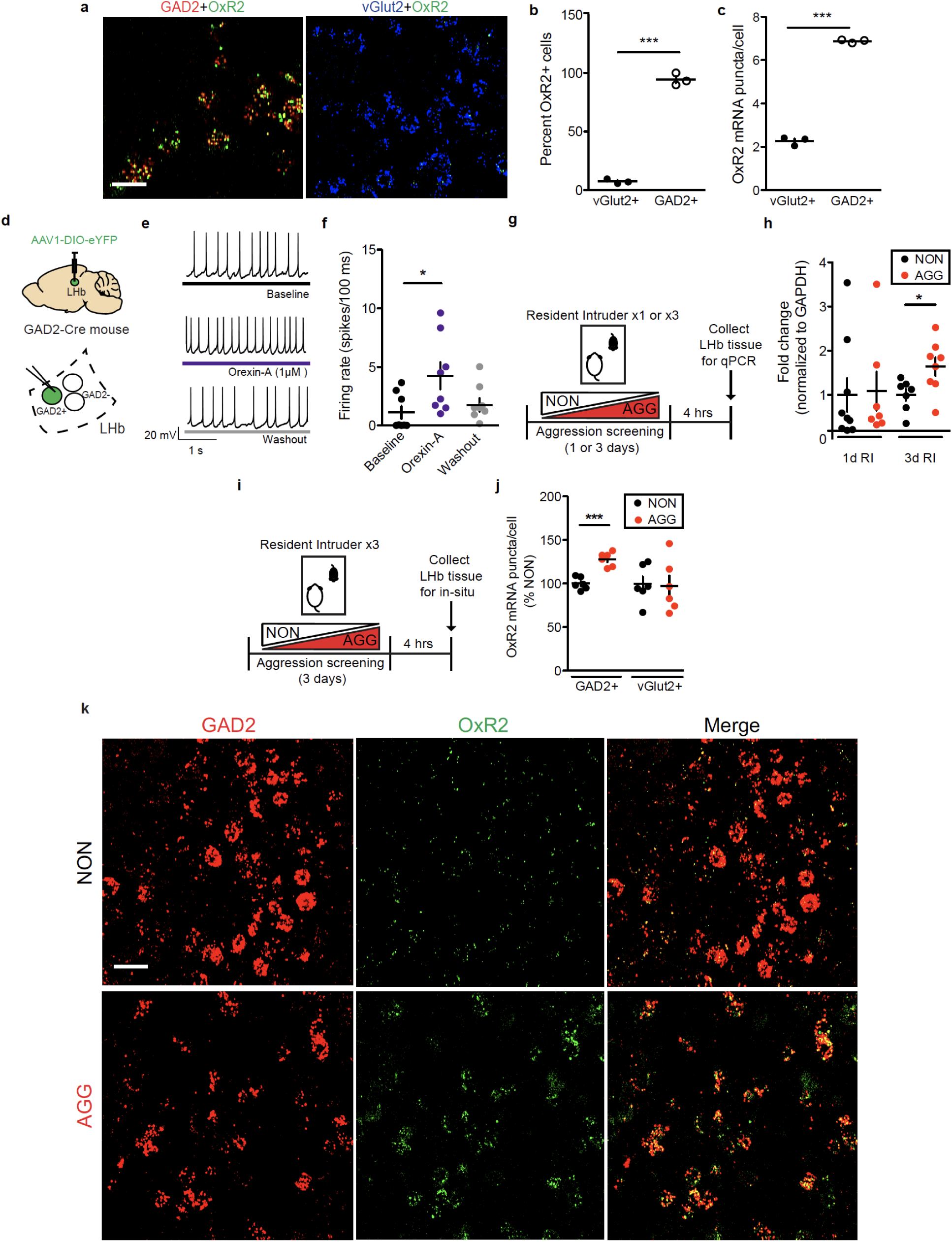
Characterization of an LHb orexin circuit. **a,** Representative *in-situ* hybridization (ISH) images showing OxR2 expression in GAD2 and vGlut2 LHb neurons, scale bar=20 μm. **b,** OxR2 is expressed primarily in GAD2 neurons compared to vGlut2 neurons (student’s t-test, n=3 animals, 1-2 slices per animal, t(4)=15.67, p<0.0001). **c,** GAD2 neurons express more OxR2 mRNA than vGlut2 neurons (student’s t-test, n=3 animals, 1-2 slices per animal, t(4)=15.67, p<0.0001). **d,** Surgical manipulations and experimental schematic for orexin bath application experiments. **e,** Representative traces from a GFP-positive neuron (GAD2 neuron) during baseline (top), orexin-A (middle), and washout (bottom) conditions. **f,** Orexin bath application increased the firing rate of GFP-positive neurons (Kruskal-Wallis one-way ANOVA with repeated measures, n=6 mice, n=9 cells, Kruskall-Wallis statistic=7.115, p=0.0285, Dunn’s multiple comparisons baseline vs. orexin-A, p<0.05). **g,** Experimental timeline for LHb qPCR experiments. **h,** Following 3 days of RI, AGGs expressed significantly more LHb OxR2 mRNA than NONs (3 days RI: student’s t-test, n=7 NONs, n=8 AGGs, t(13)=2.496, p=0.0268; 1 day RI: student’s t-test, n=9 NONs, n=7 AGGs, t(14)=0.1433, p=0.881). **i,** Experimental timeline for OxR2 ISH experiments. **j,** Following 3 days of RI, AGGs displayed increased GAD2 neuron OxR2 mRNA compared to NONs (student’s t-test, n=5 NONs, n=6 AGGs, GAD2 neurons: t(9)6.039, p=0.0002; vGlut2 neurons: t(9)=0.6735, p=0.5175). **k,** Representative images of ISH for OxR2, GAD2, in AGGs and NONs following 3 days of RI, scale bar=20 μm. *p<0.05, ***p<0.001 All data are expressed as mean ± SEM.

To investigate whether LHb OxR2 signaling is implicated in aggression, we screened mice in RI, categorized them as either AGGs or NONs, and collected LHb tissue to perform qPCR for OxR2. We found that following three days of RI, AGGs displayed increased OxR2 expression compared to NONs (Fig. 5g-h, S8). We did not observe this increase following one day of RI, indicating that LHb OxR2 expression increases as a consequence of repeated aggression in RI. Next, we used ISH to label OxR2 mRNA in LHb GAD2 and vGlut2 neurons in AGGs and NONs following 3 days of RI. This was necessary to verify that OxR2 expression increased in GAD2 LHb neurons and to control for the possibility of extra-LHb tissue contamination of tissue punches in our qPCR experiments. As expected, we detected increased OxR2 expression in AGGs compared to NONs specifically in GAD2 neurons with no change in vGlut2 neuron OxR2 expression (Fig. 5i-k, S8). These data provide support for a model whereby increased orexin signaling in LHb GAD2 neurons as a consequence of repeated aggression experience promotes aggression and its rewarding properties, possibly through heightened local inhibition of LHb principal neurons.

### Manipulation of the LHb orexin circuit

In order to verify the *in-vivo* effects of orexin on the activation of GAD2 and vGlut2 neurons in the LHb, we optogenetically stimulated orexin terminals in the LHb of orexin-Cre mice injected with AAV-DIO-ChR2 in the LH and performed ISH for Fos and GAD2 or vGlut2 (Fig. 6a). We found that optogenetic stimulation of orexin neurons in the LHb (20 Hz, 20 ms pulses, 7mW for 30 minutes) reduced total LHb Fos expression as well as Fos expression in vGlut2 positive LHb neurons, but increased Fos expression in GAD2 neurons compared to mice that received the AAV-DIO-eYFP control virus (Fig. 6b-d). Consistent with a decrease in overall Fos expression observed in the LHb following orexin input stimulation, we also found that systemic antagonism of OxR2 with the selective antagonist N-Ethyl-2-[(6-methoxy-3-pyridinyl)[(2-methylphenyl)sulfonyl]amino]-N-(3-pyridinylmethyl)-acetamide (EMPA; 30 mg/kg, intraperitoneal) increased the activity of the LHb as measured by fiber photometry in mice expressing a non-conditional AAV-GCaMP6 in the LHb (Fig. S9). Moreover, systemic antagonism of OxR2 with EMPA (30 mg/kg, intraperitoneal) also reduced aggression and aggression CPP without affecting locomotion or anxiety-like behavior (Fig. S9). Together, these results provide further support for our model and suggest that locally-inhibitory LHb GAD2 neurons are activated by orexin originating from the LH.

**Figure 6:**
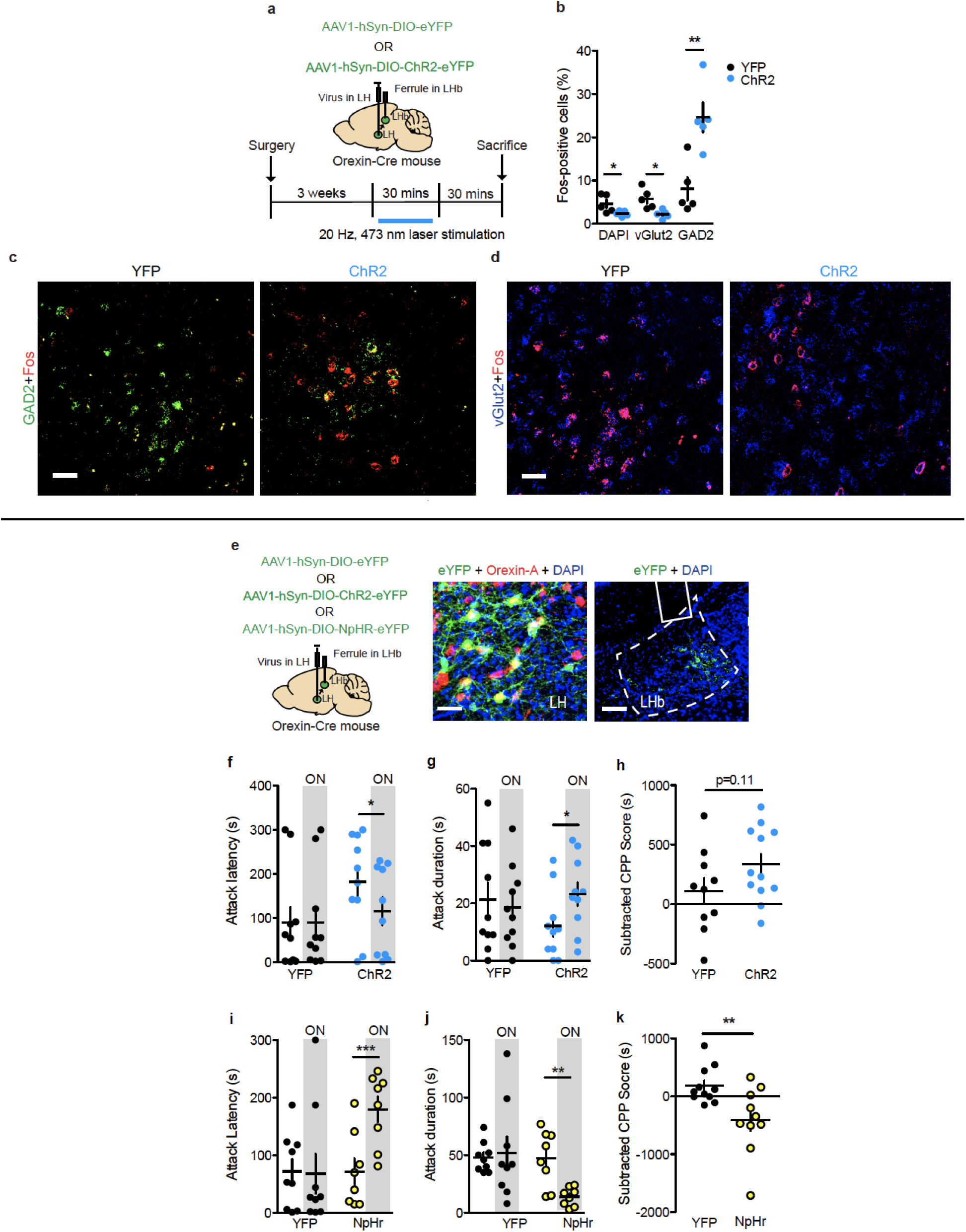
Optogenetic manipulation of orexin inputs to the LHb modulates aggressive behavior. **a,** Surgical manipulations and experimental timeline for optogenetic stimulation of orexin terminals in the LHb followed by *in-situ* hybridization (ISH) for Fos, GAD2, and vGlut2. **b,** Optogenetic stimulation of orexin terminals in the LHb increased Fos in GAD2 neurons and decreased Fos in vGlut2 neurons and DAPI positive cells (student’s t-test, n=5 per group; GAD2: t(8)=3.854, p=0.0048; vGlut2: t(8)=3.236, p=0.0120; DAPI: t(8)=2.340, p=0.0475). **c,** Representative images of Fos expression in LHb GAD2 neurons from YFP and ChR2 mice, scale bar=40 μm. **d,** Representative images of Fos expression in LHb vGlut2 neurons from YFP and ChR2 mice, scale bar=40 μm. **e,** Surgical manipulations and representative viral infection images for optogenetic manipulation of orexin terminals in the LHb, scale bar=150 μm. **f,** Optogenetic stimulation of orexin terminals reduced attack latency in RI (NpHr, paired t-test, n=10 mice, t(9)=2.354, p=0.043). **g,** Optogenetic stimulation of LHb orexin terminals increased attack duration in RI (NpHr, paired t-test, n=10 mice, t(9)=2.335, p=0.0444). **h,** Optogenetic stimulation of LHb orexin terminals did not significantly increase aggression CPP (student’s t-test, n=10 mice per group, t(18)=1.642, p=0.1140). **i,** Optogenetic inhibition of LHb orexin terminals increased attack latency in RI (NpHr, paired t-test, n=8 mice, t(7)=6.671, p=0.0003). **j,** Optogenetic inhibition of LHb orexin terminals reduced attack duration in RI (NpHr, paired t-test, n=8 mice, t(7)=4.252, p=0.0038). **k,** Optogenetic inhibition of LHb orexin terminals reduced aggression CPP (student’s t-test, n=11 YFP, n=10 NpHr, t(19)=2.993, p-0.0085). * p<0.05, ** p<0.01, ***p<0.001. All data are expressed as mean ± SEM.

Next, we tested whether orexin inputs to the LHb play a functional role in aggression and aggression reward. To do this, we injected AAV-DIO-ChR2 into the LH of orexin-Cre mice and implanted a fiber above the LHb (Fig. 6e). Optogenetic stimulation of orexin terminals in the LHb (20 Hz, 20 ms pulses, 7mW) promoted aggression and modestly increased aggression CPP in AGGs (Fig. 6f-h). However, optogenetic stimulation of orexin inputs to the LHb was not sufficient to initiate aggression in NONs (Fig. S5). To confirm that these behavioral effects in AGGs were mediated by OxR2, we performed optogenetic stimulation of LHb orexin terminals in mice expressing a microRNA against OxR2 (AAV-miR-OxR2) in the LHb. Stimulation of LHb orexin terminals under these conditions did not alter aggressive behavior (Fig. S10, S11). To determine whether inhibition of orexin terminals in the LHb reduces aggressive behavior, we injected AAV-DIO-NpHr in the LH of orexin-Cre mice and implanted a fiber above the LHb. Optogenetic inhibition of LHb orexin terminals in AGGs indeed reduced aggression and aggression CPP (Fig. 6i-k) but did not affect aggression in NONs (Fig. S5). Together, these data indicate that orexin signaling in the LHb modulates aggression as well as its rewarding properties via an OxR2-dependent mechanism.

To further verify the necessity of OxR2 signaling in LHb GAD2 neurons in aggression, we performed cell-type specific knockdown of OxR2 in GAD2 neurons using a microRNA-based approach (Fig. 7a-b, S11). Knockdown of OxR2 in LHb GAD2 neurons reduced aggression and aggression CPP (Fig. 4m-o), recapitulating the behavioral effects of direct NpHR-mediated inhibition of GAD2 LHb neurons (Fig. 7c-e). Importantly, knockdown of OxR2 in GAD2 neurons did not alter locomotor behavior or time spent in the center of an open field, indicating that effects of this manipulation on aggression were not mediated by changes in overall activity or anxiety (Fig. S13). Consistent with our previous findings over-expression of OxR2 in LHb GAD2 neurons was insufficient to promote aggressive behavior in NONs (Fig. S5, S12). These data support the conclusion that OxR2 signaling, particularly in LHb GAD2 neurons, is involved in regulating the reinforcing properties of aggression, but not its initiation.

**Figure 7:**
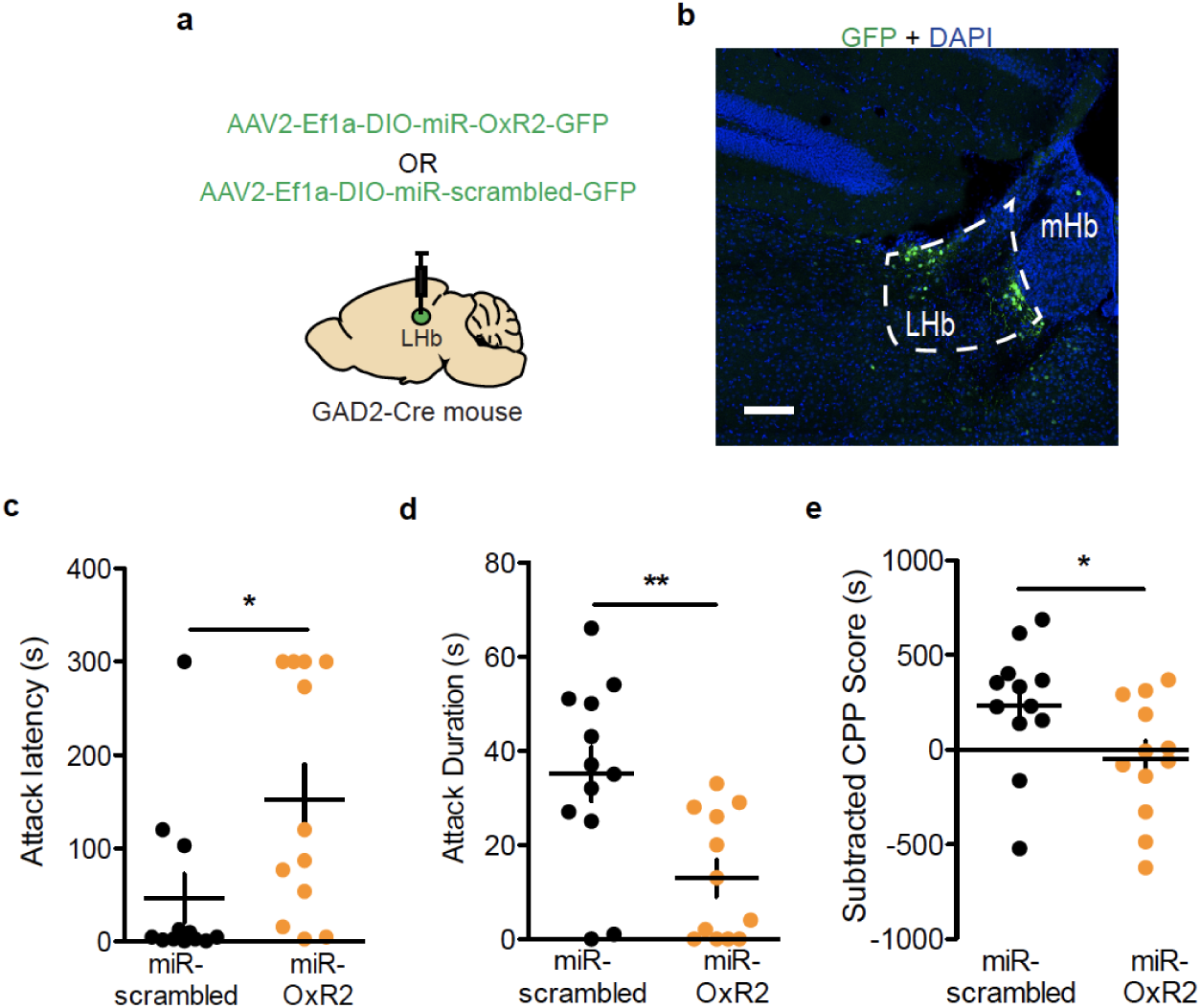
Knockdown of OxR2 in GAD2 LHb modulates aggressive behavior. **a,** Surgical manipulations for GAD2 neuron-specific knockdown of OxR2 in the LHb. **b,** Representative viral infection for GAD2 neuron-specific knockdown of OxR2 in the LHb, scale bar=300 μm. **c,** OxR2 knockdown in GAD2 LHb neurons increased attack latency in RI (student’s t-test, n=12 per group, t(22)=2.322, p=0.0299). **d,** OxR2 knockdown in GAD2 LHb neurons reduced attack duration in RI (student’s t-test, n=12 per group, t(22)=3.183, p=0.0043). **e,** OxR2 knockdown in GAD2 LHb neurons reduced aggression CPP (student’s t-test, n=12 per group, t(22)=2.155, p=0,0424). * p<0.05, **p<0.01. All data are expressed as mean ± SEM.

## Discussion

While interpersonal violence negatively affects the physical and mental wellbeing of a vast number of individuals worldwide, our current understanding of the neural circuitry and molecular mechanisms driving heightened aggression remains tenuous. Here, we used a multidisciplinary approach to monitor, manipulate, and functionally map a previously uncharacterized inhibitory cell type in the LHb that may influence the expression of aggression by altering its rewarding properties. This cell type is under crucial modulatory control by the neuropeptide orexin, which may influence its capacity to promote aggression and a preference for aggression-paired contexts.

Our results using *in vivo* fiber photometry to measure neural activity provide important new information regarding the dynamics of LHb activity during aggression and highlight adaptations occurring in LHb neurons as a result of previous fight experience. We demonstrate here that attacks are time-locked to reductions in overall LHb activity, which agrees with previous studies showing that functional inhibition of LHb neurons can enhance aggression and its rewarding properties ^14, 20^. Furthermore, we found that as AGGs learn to associate particular contexts with aggressive social interactions involving submissive intruders, the LHb displays reduced activity in these contexts even in the absence of the intruder, suggesting the structure encodes information about the rewarding properties of aggression and aggression-paired contexts based on previous experience. These findings are consistent with recent reports describing plasticity in LHb neurons as animals learn to associate cues predicting aversive or rewarding stimuli^36, 37^. While the LHb as a whole may reduce its activity during aggression or in response to aggression-related cues, we have identified a small subset of GAD2-expressing LHb neurons are robustly activated on a similar time scale to the whole LHb, suggesting that the LHb integrates the activity of multiple local cell types to promote output patterns that modulate aggression and its rewarding properties.

Our study provides the first evidence that GAD2 neurons in the LHb are capable of producing local inhibition to suppress the activity of principal neurons within the LHb that, when active, typically encode aversion. This finding brings into question the currently accepted model of LHb circuit dynamics and connectivity by adding an element of inhibitory network regulation within the LHb that was previously uncharacterized. However, many questions about these neurons remain. First, do LHb GAD2 neurons release neuropeptides or other neurotransmitters besides GABA? Notably, we found no evidence of excitatory (glutamatergic) currents in LHb principal neurons elicited from GAD2 neuron stimulation (Fig 4a-b), but this does not preclude the possibility that other messengers are also released from these neurons. In fact, a very recent study that performed single-cell sequencing of LHb neurons found that cells expressing GAD2 may also express transcripts encoding precursors for neuropeptides such as Pituitary adenylate cyclase-activating polypeptide and enkephalin^23^. Future studies should aim to verify co-expression of GAD2 and these neuropeptide precursors using *in-situ* hybridization. Second, are LHb GAD2 neurons able to exert broad inhibitory control over LHb projection neurons in aggression, or is projection-specific connectivity important? Notably, two major outputs of the LHb—the dorsal raphe nucleus (DRN) and ventral tegmental area (VTA)—have been previously implicated in aggressive behavior^38, 39^, but the role of each of these target regions in controlling motivational aspects of aggression like preference for aggression-paired contexts remains poorly understood. Future investigations should attempt to map the intra- and inter-LHb targets of GAD2 neurons that are important for regulating aggression. Third, what role, if any, do other inputs to LHb GAD2 neurons play in aggression? In addition to expressing OxR2, LHb GAD2 neurons have also been shown to express estrogen and vasopressin receptors^25^, both of which have been implicated previously in aggression^40, 41^. It is also possible that aggression-related nuclei projecting to the LHb, like the prefrontal cortex^42^ and the lateral septum^43^, target LHb GAD2 neurons directly to regulate aggression, and this should also be tested in future studies.

Our results suggest that in the context of appetitive aggression, LHb GAD2 neurons both encode reward-related signals at the completion of attacks and promote future attacks. This is not unlike previous reports that VTA dopamine neurons both encode reward-related signals following the initiation of prosocial interactions and promote future prosocial interactions^44^. The fact that optogenetic stimulation of LHb GAD2 neurons enhanced aggression in AGGs but did not elicit aggression in NONs suggests that LHb GAD2 neurons are not mediating attack initiation in this context. However, we know from our CPP studies that LHb GAD2 neurons enhance seeking behavior for aggression-associated contexts. Thus, we propose that optogenetic stimulation of LHb GAD2 neurons during RI enhances the perceived reward magnitude of each attack such that the drive to carry out subsequent attacks is enhanced through repeated positive reinforcement. This may ultimately involve feedback circuits between reward centers downstream of the LHb like the VTA and aggression initiation centers like the ventromedial hypothalamus (VMH) and medial amygdala. For example, by providing local inhibition to LHb principal neurons, GAD2 neurons may effectively disinhibit the VTA, thus encoding signals to pre-empt future aggressive behavior.

Though our data are consistent with a wide range of studies demonstrating a role for orexin in motivated behaviors, orexin has not previously been implicated in aggression. For example, a number of groups have shown that inhibition of orexin signaling reduces CPP, self-administration, and relapse for drugs of abuse^29, 32, 33, 45–47^. In addition, orexin neurons are activated by cues and contexts associated with natural rewards like food^48, 49^ and sex^50^. We were surprised, therefore, to find activation of LHb GAD2 neurons did not potentiate – or disrupt – the preference displayed during the testing phase of our palatable food CPP assay. While projections from LH glutamate neurons to the LHb indeed seem to impact feeding behaviors^51^, it seems likely that non-orexinergic glutamate neurons drive this effect. Instead, LH orexin neurons have been shown to influence feeding behavior through projections to regions outside the habenula, including the VTA^29^. Thus, it is possible these projections are insufficient to influence feeding through GAD2 LHb cells, instead acting through principal neurons in this region to regulate food intake. While orexin neurons are relatively few in number, their projections beyond the LHb are particularly enriched in regions like the DRN^38^, VTA^39^, and basal forebrain^14^, all of which play vital roles in both motivation and aggression. Notably, orexin neurons receive inputs from neurons in the ventromedial hypothalamus (VMH), a region shown to be essential for initiating inter-male attack behavior in mice^48, 52^. This connectivity pattern positions the LH-LHb orexin neural projection as a potential link between hypothalamic attack centers and circuits controlling motivation. Thus, it will be important in future studies to investigate whether additional outputs of orexin neurons are important for controlling aggression and its rewarding properties, and whether the other known orexin receptor, OxR1, plays any role.

In addition to demonstrating that cell-type specific knockdown of OxR2 reduces aggression and preference for aggression-paired contexts without affecting locomotor behavior, we also found that systemic antagonism of OxR2 with 30 mg/kg (I.P.) EMPA reduces aggression and preference for aggression-paired contexts without affecting locomotor behavior or anxiety-like behavior in the open field test. The lack of an effect of EMPA on locomotion is somewhat surprising, since decades of research have shown that orexin signaling is integral for promoting wakefulness. Indeed, brain-wide inhibition of orexin neurons^53^ or deletion of OxR2^54^ results in narcolepsy-like phenotypes in rodents. In addition, the dual orexin receptor antagonist suvorexant is currently an FDA-approved treatment for insomnia, as it aids in the transition from wakefulness to sleep in humans^55^. While previous studies have shown that 30 mg/kg EMPA reduces spontaneous locomotion in rats^56^, this dose of EMPA does not affect spontaneous locomotion in mice^57^. Together with our study, these data may suggest that if given at sufficiently low doses, EMPA could reduce aggression without side effects on arousal. However, given that spontaneous locomotor activity is not a particularly sensitive assay for arousal, this possibility must be examined in detail in future studies.

It is important to note that these experiments were performed during the light cycle, a period of lower activity levels for nocturnal rodents such as mice. While the mechanism is not fully elucidated, aggressive behaviors do show variability across the light-dark cycle^58^. Circadian rhythms may also crucially affect the LHb in particular, which may subsequently exert varying effects throughout the light-dark cycle on downstream targets regulating a variety of motivated behaviors^59^. It is therefore possible these findings may differ had experiments been performed during the dark cycle instead of the light cycle, so caution must be taken when interpreting these results. The ramifications of the dark cycle on aggressive behaviors – and activities of this putative orexin LH to LHb GAD2 pathway – should be investigated more closely in follow-up studies.

In conclusion, we illustrate here that orexin likely plays an important role in modulating aggression and preference for aggression-paired contexts through the activation of inhibitory LHb GAD2 neurons. These findings may have important implications for the treatment of aggression in psychiatric patients.

## Online Methods

### Animals

For experiments in wild-type animals, 4-month old male CD-1 (ICR) mice (sexually experienced retired breeders; Charles River Laboratories) were used as subjects. Subjects were confirmed by CRL to have equal access, experience, and success as breeders. For experiments in transgenic animals, C57BL/6J heterozygous Orexin-cre-IRES-GFP (gift from A. Yamanaka, see ^27^) or homozygous GAD2-Cre (Jackson Laboratory) mice were crossed to wild-type CD-1 mice and the F1 generation was used for experiments. This strategy was necessary to ensure a wide range of aggressive phenotypes in experimental animals, as C57BL/6J mice display relatively low levels of aggressive behavior ^13^. At 3 months of age, F1 transgenic male mice were paired with F1 female mice for two weeks to gain sexual experience before being utilized for experiments. For experiments in transgenic mice, male littermates were randomly assigned to experimental groups. 8-9 week male C57BL/6J mice (20-30g; The Jackson Laboratory) were used as novel intruders. All mice were allowed one week of acclimation to the housing facilities before the start of experiments. Wild-type CD-1 and F1 transgenic mice were singly housed and C57BL/6J mouse were housed in groups of 5. All mice were maintained on a 12 h light:dark cycle (7AM to 7PM) with *ad libitum* access to food and water. Consistent with many previous studies on aggression^9, 39, 58, 60^, behavioral experiments were conducted during the light phase. However, it is unclear whether our results are directly comparable to studies performed during the dark phase. Procedures were performed in accordance with the National Institutes of Health Guide for Care and Use of Laboratory Animals and the Icahn School of Medicine at Mount Sinai Institutional Animal Care and Use Committee.

### Aggression screening/Resident Intruder (RI) test

Aggression screening was performed as previously described by utilizing the resident intruder (RI) test ^14^. After a minimum of one week of habituation to home cages, experimental mice were exposed to a novel C57BL/6J intruder for 5 min daily over 3 consecutive days. Each intruder presentation was performed in the home cage of the experimental mouse between 12-3 PM daily under white light conditions. During RI sessions the cage top was removed to allow for unobstructed viewing and video recording of sessions. The duration and number of screening sessions were selected to prevent induction of stress- and anxiety-related behaviors in experimental CD-1 or F1 hybrid mice^14^. All RI sessions were video recorded with a digital color video camera. Two blind observers recorded (1) the latency to initial aggression and (2) the total duration of aggression. The initiation of aggression was defined by the first clear physical antagonistic interaction initiated by the resident mouse (usually a bite), not including grooming or pursuit behavior. Aggression was considered completed when the resident mouse had reoriented away from the intruder following the initiation of attack. This definition allows for slight breaks (less than 5 s) in continuous physical interaction within an aggressive bout, assuming the resident mouse has remained oriented towards the intruder throughout. Resident mice were defined as AGGs if they initiated aggression during all three screening sessions, while NONs were defined as those that showed no aggression during any screening sessions. Aggression screening was halted if an intruder showed any signs of injury in accordance with our previously published protocols^14, 61^.

### Aggression conditioned place preference (CPP)

We carried out the aggression CPP protocol according to Golden et al.^14^. Briefly, this task consisted of three phases: pre-test, acquisition (conditioning), and test. Mice were acclimated to the testing facility for 1 h before all testing. All phases were conducted under red light and sound-attenuated conditions. The CPP apparatus (Med Associates) consisted of two unique conditioning chambers with a neutral middle zone that allowed for unbiased entry into either conditioning chamber at the initiation of each trial. During the pre-test phase, mice were placed into the middle chamber of the conditioning apparatus and allowed to freely explore the apparatus for 20 min. There were no group differences in bias for either chamber, and conditioning groups were balanced in an unbiased fashion to account for pre-test preference. The acquisition phase consisted of three consecutive days with two conditioning trials each day for a total of 6 acquisition trials. Morning trials (between 8-10 AM) and afternoon trials (between 3-5 PM) consisted of experimental mice confined to one chamber for 10 min while in the presence or absence of a novel C57BL/6J intruder mouse. All groups were counterbalanced for the conditioning chamber. A total of 3 conditioning trials to the intruder-paired and intruder-unpaired context were performed. On the test day, experimental mice were placed into the middle arena without any intruders and allowed to freely explore the apparatus for 20 min. For optogenetic experiments, light was delivered during the full duration of the test phase only. Total locomotor activity was also recorded to ensure equal exploratory behavior between groups. Behavioral analysis of aggression CPP was performed by calculating (1) CPP score (test phase duration in paired chamber minus test phase duration in unpaired chamber) (2) subtracted CPP score (test phase duration in paired chamber subtracted by pre-test phase duration in paired chamber).

### Open field test

The open field test was performed as previously described ^62^. One week after the last RI, experimental mice were acclimated to the testing facility for 1 h before testing. Open-field tests were performed in black plexiglass arenas (42 x 42 x 42 cm; Nationwide Plastics) under red light conditions. Testing sessions lasted for either 5 minutes (GAD2-specific OxR2 knockdown and systemic EMPA experiments) or 10 minutes (non-conditional OxR2 knockdown experiment). Behavior was tracked with Noldus Ethovision (Noldus Interactive Technologies) to record the total distance moved, time spent in the entire arena, and time spent in the delineated “center zone” or “corner zones” of the arena (24 x 24 cm).

### Behavioral pharmacology

Mice were injected intraperitoneally with either 30 mg/kg EMPA (Tocris; 4558) dissolved in 0.3% v/v Tween-80 in saline or vehicle (0.3% v/v Tween-80 in saline alone) 25 minutes prior to behavioral testing. For RI tests, animals were given EMPA one day and vehicle the other (counterbalanced for order, within-subjects design). For CPP and open field tests, half of the animals were given EMPA and half were given vehicle (between-subjects design).

### Perfusion and brain tissue processing

For immunohistochemistry and histology, mice were given a lethal dose of 15% chloral hydrate and transcardially perfused with cold PBS (pH 7.4) followed by fixation with cold 4% paraformaldehyde (PFA) in PBS. Brains were dissected and post-fixed for 24 h in 4% PFA. Coronal sections were prepared on a vibratome (Leica) at 50 μm to assess viral placement and perform immunohistochemistry.

For *in-situ* hybridization, mice were rapidly decapitated and brains were removed and flash frozen in −30 degrees C isopentane for 30 s and then stored at −80 degrees C until sectioning. Coronal sections for *in-situ* were prepared on a cryostat at 16 μm thickness and mounted directly on slides.

For real-time quantitative PCR (RTqPCR), mice were rapidly decapitated and the brains were extracted and placed in ice-cold PBS. Bilateral LHb 1 mm diameter, 1 mm thick, tissue punches were taken and immediately flash frozen on dry ice and stored at −80 degrees until RNA extraction.

### RNA extraction, generation of cDNA, and RTqPCR

RNA was isolated from either brain tissue or HEK293 cells using TRIzol (Invitrogen) and chloroform phase separation. The clear RNA layer was processed with the RNAeasy MicroKit (Qiagen), analyzed with the NanoDrop (Thermo Fisher Scientific), and 500 ng of RNA was reverse transcribed to cDNA with qScript (95048-500; Quanta Biosciences). The resulting cDNA was diluted to 1 ng/μl.

For qPCR of OxR1 and OxR2, 3 μl of cDNA was combined with 5 μl of Perfecta SYBR Green (95054-02K; Quanta Biosciences), forward/reverse primers (1 μl total), and 1 μl of water. Samples were heated to 95 degrees C for 2 min followed by 40 cycles of 95 degrees C for 15s, 60 degrees C for 33 s, and 72 degrees C for 33 s. Analysis was performed using the delta_deltaC(t) method. Samples were normalized to GAPDH. Primers used were as follows: GAPDH (F: AAC GGC ATT GTG GAA GG, R:GGA TGC AGG GAT GAT GTT CT), orexin receptor 1 (OxR1) (F: ATC CAC CCA CTG TTG TT, R: GGC CAG GTA GGT GAC AAT GA), orexin receptor 2 (OxR2) (F: CAT CGT TGT CAT CTG GAT CG, R: GGC ACC AGA GTT TAC GGA AT).

For qPCR of all other transcripts, reactions were performed with 30 ng of cDNA per 10ul reaction, Taqman Fast Advanced Master Mix (Life Technologies), and Taqman probes (Life Technologies) on a QuantStudio 7 Flex Real-Time PCR system (Life Technologies). Taqman probes used were: *Avpr2 (*Mm01193534_g1), *Drd1*(Mm02620146_s1), *Drd4* (Mm00432893_m1), *Glp1r* (Mm00445292_m1), *Htr3a* (Mm00442874_m1), *Mchr1* (Mm00653044_m1), and *Nmur* (Mm00515885_m1). Samples were held at 95 degrees C for 20s, then cycled from 95 degrees C for 1s to 60 degrees C for 20s for 40 cycles. Gene expression was normalized to housekeeping genes *Abt1* (Mm00803824_m1) and *Hprt* (Mm03024075_m1) using the delta_deltaC(t) method.

### Generation and validation of AAV2 Cre-dependent OxR2 viral constructs

To create an effective Cre-dependent knockdown virus for OxR2, we utilized a micro-RNA (miR) based approach. The miR was bicistronically expressed with IRES-eGFP for simple identification of infected cells. Briefly, we generated the OxR2 miR using the BLOCK-iT ™ Pol II miR RNAi expression vector kit (Thermo Fisher Scientific). We used the shRNA sequence from our non-conditional OxR2 virus, which was previously validated to inhibit OxR2 expression ^63^. A scrambled sequence was used as the control. We inserted the miR-containing oligonucleotides into a pcDNA6.2-GW-miR vector provided by the kit. The miR sequences, along with the 5’ and 3’ flanking regions, were then sub-cloned into a bicistronic IRES-GFP vector (pAAV-IRES-GFP, Cell Biolabs). This vector was non-conditional and can be expressed in mammalian cells. Suppression of OxR2 with the miR construct was validated in N2A cells by qPCR (see above). Once validated *in-vitro*, the miR-IRES-GFP sequence was inserted into a Cre-dependent AAV2.Ef1a.DIO vector and packaged (Virovek Inc.) to produce AAV2-Ef1a-DIO-miROxR2-IRES-GFP-SV40pA and pAAV-Ef1a-DIO-miRscrambled-IRES-GFP-SV40pA. For over-expression of OxR2, we purchased a non-conditional DNA construct containing OxR2 (pcDNA-CMV-OxR2-Myc-FLAG-hGH-SV40, Genecopoeia) and verified over-expression in N2A cells using qPCR. To create an effective Cre-dependent over-expression virus for OxR2, the sequence for OxR2 was inserted into a Cre-dependent AAV2.Ef1a.DIO vector and packaged to produce AAV2-Ef1a-DIO-OxR2-SV40pA (Virovek, Inc.). These AAVs were subsequently validated *in-vivo* through injection into GAD2-cre mice and *in-situ* hybridization for GFP, GAD2, and OxR2.

### Immunohistochemistry (IHC), *in-situ* hybridization (ISH), and confocal microscopy

For IHC experiments, sections were incubated overnight in blocking solution (3% normal donkey serum, 0.3% Triton X-100 in PBS), washed three times in PBS for 10 min (30 min total), then incubated for 24 h in primary antibodies diluted in blocking solution (goat anti-Orexin-A (Santa Cruz Biotechnology, sc-8070;1:500); chicken anti-GFP (Aves Labs, GFP-1020; 1:1000). Sections were then washed three times in PBS for 10 min (30 min total), incubated for 2 h in secondary antibodies diluted in blocking solution (donkey anti-goat Cy3 1:400 (Jackson ImmunoResearch; 705-165-003); donkey anti-chicken AlexaFluor 488 1:400 (Jackson ImmunoResearch; 703-545-155), and washed three times in PBS for 10 min (30 min total). Finally, sections were counterstained with 1 ug/ml DAPI (Sigma) for 10 min and mounted on slides. Sections were allowed to dry on slides overnight, dehydrated with ethanol and then Citrisolv, and cover-slipped with ProLong Diamond Antifade Mountant (Invitrogen; P36970). IHC of orexin axons and GAD2 LHb cells were imaged using the AiryScan method on a LSM 880 confocal microscope (Carl Zeiss) and analyzed with Imaris (Bitplane). All other IHC was imaged with a LSM 780 confocal microscope (Carl Zeiss) and analyzed with FIJI (ImageJ) software.

For ISH experiments, we utilized the RNAScope Multiplex Fluorescent *in-situ* kit (Advanced Cell Diagnostics) according to the manufacturer’s instructions. Briefly, fresh frozen sections were fixed in ice-cold 4% PFA in PBS for 15 minutes, serially dehydrated with EtOH (50%, 75%, 100%; each for 2 minutes), and pretreated with a protease (Protease IV, RNAScope) for 30 min. Proprietary probes for eGFP, glutamic acid decarboxylase 2 (GAD2), or vesicular glutamate transporter 2 (vGlut2), or orexin receptor 2 (OxR2) (Advanced Cell Diagnostics) were hybridized at 40 degrees C for 2 h, serially amplified, counterstained with 1 ug/ml DAPI for 2 min, and immediately cover-slipped with EcoMount mountant (Biocare Medical). *In-situ* hybridization was imaged with a Zeiss LSM 780 confocal microscope at 20x, 40x, or 63x magnification. mRNA puncta were quantified manually, blinded to experimental condition, using FIJI software (Image J).

### *In-vitro* electrophysiology

#### Slice preparation

Adult male mice were anesthetized with isoflurane, decapitated, and the brain was immediately removed and submerged into ice-cold sucrose-artificial CSF (aCSF) comprising (in mM) 233.7 sucrose, 26 NaHCO_3_, 3 KCl, 8 MgCl_2_, 0.5 CaCl_2_, 20 glucose, and 0.4 ascorbic acid. Coronal sections (350 microns thick) were sliced using a Leica VT1000S vibratome and allowed to equilibrate in recording aCSF (in mM: 117 NaCl, 4.7 KCl, 1.2 MgSO_4_, 2.5 CaCl_2_, 1.2 NaH_2_PO_4_, 24.9 NaHCO_3_, and 11.5 glucose) oxygenated with 95% O_2_-5% CO_2_ for 1 hour at RT. Slices were then transferred to the recording chamber where they were maintained at 31 degC and perfused (1.5 ml/min) with oxygenated aCSF.

#### Orexin bath application recordings

2 weeks prior to recording, GAD2-cre were injected in the lateral habenula (see coordinates in following section) with AAV-hSyn-DIO-eYFP (UPenn Viral Core). Mice were then returned to their home cage to allow for viral expression before being killed for slice recordings. During recording, cells were identified as GAD2-positive by expression of eYFP in the lateral habenula. Neurons were visualized on an upright epifluorescence microscope (BX50WI; Olympus) with a 40X water-immersion objective and an infrared CCD monochrome video camera (Dage-MTI). Whole-cell recordings were performed with glass micropipettes (resistance 2-4 MΩ) pulled from borosilicate glass capillaries using a P-87 micropipette puller (Sutter Instruments). The pipettes were filled with an intracellular solution containing: 124 mm K-gluconate, 10 mm HEPES, 10 mm phosphocreatine di(Tris), 0.2 mm EGTA, 4 mm Mg_2_ATP, and 0.3 mm Na_2_GTP, adjusted to an osmolarity of 280-290 mOsm and pH of 7.3. Recordings were made with a Multiclamp 700B (Molecular Devices) in current-clamp mode. Analog signals were low-pass filtered at 2 kHz and digitized at 5kHz using a Digidata 1440A interface and pClamp10 software (Molecular Devices). Gigaseal and access to the intracellular neuronal compartment was achieved in voltage-clamp mode, with the holding potential set at −70 mV. After rupturing the membrane, intracellular neuronal fluid equilibrated with the pipette solution without significant changes in series resistance or membrane capacitance. Cells were allowed to normalize for 5 minutes before recording. To test response of cells to orexin A (Phoenix Pharmaceuticals), we applied orexin A (1 µM) for 5 minutes in the circulating bath after a 2-minute baseline period, followed by washout for 45-60 minutes. Orexin A was prepared freshly and dissolved in aCSF before being added to recording bath.

Offline analysis was performed using Clampfit (Molecular Devices). Spikes were counted during baseline, orexin wash-in, and wash-out periods. To account for bath circulation, spikes were counted 2 minutes after drug was added or after wash-out began. Kruskal-Wallis test with Dunn’s post-hoc tests for multiple comparisons was conducted to analyze mean firing rate between groups (baseline, orexin wash-in, and wash-out) using GraphPad Software Prism (version 5.01).

#### Slice optogenetic stimulation of LHb GAD2 neurons recording

2 weeks prior to recording, GAD2-Cre mice were injected with AAV1-DIO-ChR2-eYFP (UPenn Viral Core) in the LHb. Mice were then returned to their home cage to allow for viral expression before being killed for slice recordings. Recordings were obtained with borosilicate glass electrodes (5-8 MΩ resistance) filled with voltage clamp internal solution (in mM: 120 Cs-methanesulfonate, 10 HEPES, 10 Na-phosphocreatine, 8 NaCl, 5 TEA-Cl, 4 Mg-ATP, 1 QX-314, 0.5 EGTA, and 0.4 Na-GTP). Cells were visualized on an upright DIC microscope equipped with a 460 nm objective-coupled LED (Prizmatix) for verification of ChR2 expression as well as optogenetic cellular manipulations. Data were low-pass filtered at 3 kHz and acquired at 10 kHz using Multiclamp 700B and pClamp 10 (Molecular Devices). The polarity of light-evoked stimulation (λ 460, 1 mW, 1-5 ms) of ChR2+ terminals was determined by clamping cells at −70mV (excitatory responses) and 0mV (inhibitory responses). The monosynaptic nature of light-evoked currents was confirmed by bath application of tetrodotoxin (1 μM; Abcam) and 4-aminopyridine (100 μM; Abcam) as previously described ^64, 65^. The location of cells within the LHb was confirmed visually after recording.

### Stereotaxic surgery and viral gene transfer

Mice were anesthetized with a mixture of ketamine (100 mg/kg body weight) and xylazine (10 mg/kg body weight) and placed securely in a stereotaxic frame (David Kopf Instruments). 33-gauge syringes (Hamilton Co.) were used to bilaterally infuse 0.5 μl of virus over 5 min and virus was allowed to diffuse for 5 minutes before the needle was withdrawn. For lateral hypothalamus (LH) injections, the coordinates from bregma were −1.3 mm AP, +/- 1.1 mm ML, and −5.1 mm DV at a 0-degree angle. For lateral habenula (LHb) injections, the coordinates from bregma were −1.7 mm AP, +/- 0.6 mm ML, and −2.7 mm DV at a 10-degree angle. For all optogenetics experiments animals were implanted with an optical fiber at the same time as viral injection (−2.3 mm DV). Optic fibers (Doric Lenses, MFC_200/240-0.22_2.6mm_FLT) were secured to the skull using C&B Metabond adhesive luting cement (Parkell Dental Supply). For electrophysiology experiments, GAD2-cre F1 mice were injected bilaterally in the LHb with either AAV1-hSyn-DIO-eYFP or AAV1-hSyn-DIO-ChR2.eYFP (UPenn Viral Core). For orexin optogenetics experiments, AAV1-hSyn-DIO-NpHr3.0-eYFP, AAV1-hSyn-DIO-ChR2-eYFP, or AAV1-hSyn-DIO-eYFP (UPenn Viral Core) was injected into the LH of Orexin-cre-IRES-GFP F1 mice and the optic fiber was placed in the LHb. Importantly, IRES-GFP labeling of orexin neurons is not visible without enhancement of GFP signal using immunohistochemistry, while viral eYFP is visible in perfused slices without amplification. This is evident in images of LH tissue from orexin-cre animals injected with AAV-DIO-eYFP—only a portion of the total orexin neurons are green due to infection with the virus. The specificity of viral expression in orexin positive neurons was confirmed (Fig. 5d, S5). For GAD2 neuron optogenetics experiments, GAD2-cre F1 mice were injected bilaterally in the LHb with either AAV1-hSyn.DIO-NpHr3.0-eYFP, AAV1-hSyn-DIO-ChR2-eYFP, or AAV1-hSyn-DIO-eYFP (UPenn Viral Core). For conditional knockdown of OxR2, GAD2-cre F1 mice were injected bilaterally in the LHb with either AAV2-Ef1a-DIO-miROxR2-IRES-eGFP (knockdown) or AAV2-Ef1a-DIO-eGFP (Virovek, Inc, generated for this paper). For conditional over-expression of OxR2 GAD2-Cre F1 mice were injected bilaterally in the LHb with either AAV2-EF1a-DIO-GFP or AAV2-EF1a-DIO-OxR2. For non-conditional knockdown of OxR2 during optical stimulation of orexin terminals in the LHb, orexin-Cre F1 mice were injected with AAV1-hSyn-DIO-ChR2 or AAV1-hSyn-DIO-eYFP in the LH and AAV2-hSyn-Cre plus AAV2-Ef1a-DIO-miR-OxR2-GFP or AAV2-Ef1a-DIO-miR-scrambled-GFP in the LHb. For fiber photometry in all LHb neurons, wild-type CD1 mice were injected unilaterally in the LHb with AAV1-hSyn-GCaMP6s. For fiber photometry in GAD2 neurons, GAD2-Cre mice were injected unilaterally in the LHb with AAV9-hSyn-FLEX-GCaMP6s. Optic fibers for photometry (Doric Lenses, MFC_400/430-0.48_2.7mm_MF2.5-FLT) were implanted over the LHb at the same time as viral injection and secured with dental cement. AAV1 and AAV9 viruses were allowed 2-4 weeks for expression, while AAV2 viruses were allowed 4-6 weeks for expression.

### Optogenetic stimulation

For blue light stimulation (ChR2), optical fibers (Doric Lenses, MFC_200/240-0.22_2.6mm_FLT) were connected to a 473-nm blue laser diode (Crystal Laser, BCL-473-050-M) using a patch cord with an FC/PC adapter (Doric Lenses, MFP_200/240/900-0.22_4m_FC-MF2.5). A function generator (Agilent Technologies; 33220A) was used to generate 20 ms blue light pulses at 20 Hz. The intensity of light delivered to the brain was 7-10 mW. These parameters are consistent with previously validated and published protocols^14^. In particular, 20 Hz was selected for excitation of orexin neurons since *in-vitro* experiments have demonstrated that 20 Hz stimulation is sufficient to elicit activation of orexin receptors^66^.

For yellow light stimulation (NpHR), optical fibers were connected to a 561-nm yellow laser diode (Crystal Laser, CL561-050L) using an FC/PC adapter. A function generator (Agilent Technologies; 33220A) was used to generate constant light pulses for 8 s followed by 2 s of light off. The intensity of the light delivered to the brain was 7-10 mW. These parameters are consistent with previously validated and published protocols^14^.

For all optogenetics experiments, experimental mice were habituated to patch cords for 5 days prior to testing in RI and aggression CPP. For RI experiments, mice were tested twice in the same day (in a counterbalanced fashion) in both laser on and laser off conditions with at least 4 hours between sessions (within subjects design). For CPP experiments, both groups (YFP or opsin) received laser stimulation (between subjects design).

### Real-time place preference

For real-time place-preference (RT-PP) experiments, mice were placed in the center of an open field and allowed to freely explore for 20 minutes. The time spent on each side was recorded. For the first 10 minutes of testing, one side of the open field was paired with 20 ms pulses of 20 Hz blue light stimulation (473 nm, 7-10 mW intensity). For the second 10 minutes of testing, laser stimulation was paired with the opposite side of the open field. This was done to minimize inherent bias towards one side of the open field. A RT-PP score was calculated by subtracting the total time spent in the unstimulated side from the total time spent in the stimulated side.

### Palatable food CPP

The palatable food CPP task consisted of three phases: pre-test, acquisition (conditioning), and test. Mice were acclimated to the testing facility for 1 h before all testing. All phases were conducted under red light conditions. The CPP apparatus (Med Associates) consisted of two unique conditioning chambers with a neutral middle zone that allowed for unbiased entry into either conditioning chamber at the initiation of each trial. During the pre-test phase, mice were placed into the middle chamber of the conditioning apparatus and allowed to freely explore the apparatus for 20 min. There were no group differences in bias for either chamber, and conditioning groups were balanced in an unbiased fashion to account for pre-test preference. The acquisition phase consisted of six consecutive days with two conditioning trials each day for a total of 12 acquisition trials. Morning trials (between 8-10 AM) and afternoon trials (between 3-5 PM) consisted of experimental mice confined to one chamber containing Reese’s mini peanut butter cups for 20 min and one chamber containing standard chow for 20 min. Mice were exposed to Reese’s mini peanut butter cups in the home cage prior to CPP testing to reduce effects of novelty during conditioning. All groups were counterbalanced for the conditioning chamber and conditioning time (AM vs. PM pairings with peanut butter cups). On the test day, experimental mice were placed into the middle arena and allowed to freely explore the empty apparatus for 20 min. For optogenetic experiments, light was delivered during the full duration of the test phase only and not during conditioning. Behavioral analysis of CPP was performed by calculating (1) CPP score (test phase duration in paired chamber minus test phase duration in unpaired chamber) (2) subtracted CPP score (test phase duration in paired chamber subtracted by pre-test phase duration in paired chamber).

### Fiber photometry

#### Apparatus: Resident Intruder and Aggression CPP

A fiber optic patch cord (Doric Lenses, MFP_400/430/1100-0.37_3m_FC-MF2.5) was attached to the implanted fiber optic cannula with cubic zirconia sleeves. In turn, the fiber optic cable was coupled to the apparatus for light delivery and signal measurement. GCaMP6s signal was measured by passing 490 nm LED light (Thorlabs) through a GFP excitation filter (MF469; Thorlabs) and dichroic mirrors (DMLP425, MD498; Thorlabs) into the brain and focusing emitted light onto a photodetector (2151 femtowatt receiver; Newport) after passing it back through a dichroic mirror (MD498; Thorlabs), through a GFP emission filter (MF525-39; Thorlabs), and through a 0.50 N.A. microscope lens (62-561;Edmund Optics). To account for auto-fluorescence and possible motion artifacts during testing, a second 405 nm LED not corresponding to GCaMP6s delivered light through a violet excitation filter (FB405-10; Thorlabs) and the same dichroic mirrors as the 490 nm light. This signal was similarly directed into the brain and subsequently measured with the photodetector. Light at the fiber tip ranged from 30 to 75 μW but was constant across trials over days. Simultaneous recording of both 490 and 405 nm channels was achieved through sinusoidal modulation of the LEDs at different frequencies so that the signals could be easily unmixed. Signals were collected at a rate of 381 Hz and visualized using a real-time signal processor (RX8; Tucker-Davis Technologies) and PC OpenEx software (Tucker-Davis Technologies).

#### Behavior: Resident Intruder and Aggression CPP

The timeline for aggression photometry experiments was as follows: viral injection and ferrule implantation (day 0), habituation to patch cord (days 14-15), RI recordings (days 16-18), and CPP recordings (days 21-25). Once hooked up to the apparatus, mice were allowed to rest for 10 minutes before the start of recording. Once recording was initiated, the GCaMP signal was allowed to stabilize for two minutes before the start of the behavioral trial. On each day of RI, we collected 5 minutes of baseline recordings followed by 5 minutes of intruder exposure. For CPP, photometry data was collected during the entire 20-minute pre-test and test.

#### Analysis: Resident Intruder and Aggression CPP

Analysis of signals was done using custom written MATLAB code^44, 67^, which can be obtained from the authors upon request. The bulk fluorescent signal from each channel was normalized to compare across recording sessions and animals. Change in fluorescence was calculated as a percentage of the total fluorescence signal in the GCaMP channel (deltaF/F). The 405 channel served as the control channel and was subtracted from the GCaMP channel to eliminate signals due to auto-fluorescence, bleaching, and the bending of the fiber optic cord during aggression. In general, these motion artifacts had very minimal effects on the overall GCaMP signal. Behavioral data was temporally aligned with fluorescence recording data by sending 1s-interval TTL signals to the OpenEx software from Noldus Ethovision (Noldus Interactive Technologies) behavioral recording software. To identify peak signals, we first determined the median average deviation (MAD) of the corrected/normalized data sets. Peak events that exceeded the MAD by 2.91 deviations were determined to be significant peaks, and this is in accordance with previous reports using the fiber photometry technique ^44, 67^. For analysis of LHb GCaMP activity during discrete behaviors in RI, average deltaF/F signals (%) in the two seconds before and after a discrete event (bite, approach, withdrawal) were compared. A bite was determined to occur at the moment of jaw closing on the body of the intruder, an approach at the moment of resident nose contact with any body part of the intruder, and a withdrawal at the moment body contact between resident and intruder ceased and one or both mice turned away from the site of interaction.

#### Apparatus: Palatable food consumption

A branched fiber optic patch cord (Doric Lenses, BFP_200/230/900-0.37_FC-MF1.25) connected to the fiber photometry apparatus (Neurophotometrics, Ltd.) was attached to the implanted fiber optic cannula using a cubic zirconia sleeve. To record fluorescence signals from GCaMP6s, light from a 470 nm LED was bandpass filtered, collimated, reflected by a dichroic mirror, and focused by a 20x objective. LED light was delivered at a power that resulted in 75-150 uW of 470 nm light at the tip of the patch cord. Emitted GCaMP6s fluorescence was bandpass filtered and focused on the sensor of a CCD camera. To account for auto-fluorescence and possible motion artifacts during testing, a second 560 nm LED not corresponding to GCaMP6s delivered light through a excitation filter and the same dichroic mirrors as the 470 nm light. This signal was similarly directed into the brain and subsequently measured with the CCD camera. Simultaneous recording of both 470 and 560 nm channels was achieved through an integrated camera and image splitter (Flir Blackfly S series). Signals were collected at a rate of 40 Hz and visualized using the open-source software Bonsai (http://bonsai-rx.org).

#### Behavior: Palatable food consumption

All recordings were performed in the subjects’ home cages between 10AM-12PM. The timeline for photometry experiments was as follows: viral injection and ferrule implantation (day 0), habituation to patch cord (days 14-15), then recordings with food interactions (days 21-30). For food-restricted conditions, food was removed from cages at 5PM the day before testing for a total food-restriction period of 18 hours. Once attached to the patch cord, animals habituated for a period of 10 minutes while video and LED protocols were started. Baseline signal was taken from the final 5 minutes of recording before a 5-minute exposure to food. Behaviors were scored manually using JWatcher 1.0 (Daniel T Blumenstein, Janice C Daniel, and Christopher S Evans, http://www.jwatcher.ucla.edu/).

#### Analysis: Palatable food consumption

For fiber photometry analysis, the bulk fluorescent signal from each channel was normalized to compare across recording sessions and animals. Change in fluorescence was calculated as a percentage of the total fluorescence signal in the GCaMP channel (deltaF/F). The 560 nm channel was used to control for auto-fluorescence, bleaching, and the bending of the fiber optic cord during testing. In general, these motion artifacts had very minimal effects on the overall GCaMP signal. For analysis of LHb GCaMP activity during food consumption, average deltaF/F signals in the ten seconds before and after consumption were compared.

### Statistics

All statistical details can be found in the figure legends, including type of statistical analysis used, p values, n, what n represents, degrees of freedom, and t or F values. For comparisons of two experimental groups, two-tailed paired (student’s, within subject) or unpaired (between subject) t-tests were used. For parametric data sets, comparisons of three or more groups were performed using one or two-way ANOVA tests followed by a Bonferroni posthoc test. For non-parametric data sets, comparisons of three or more groups were performed using Kruskall-Wallis one-way ANOVA followed by Dunn’s test for multiple comparisons. For all tests, p<0.05 was deemed significant. Statistical analyses were performed using Graph Pad Prism 5 software.

## Acknowledgements

The authors would like to thank Susie Feng and Carmen Ferrer for their assistance with histology, Nikos Tzavaras for his assistance with microscopy, and Virovek Inc. for cloning and packaging of AAV viruses. This work was supported by NIH grants 1R01MH114882-01 (S.J.R.), 2R01MH090264-06 (S.J.R.), P50 MH096890 and P50 AT008661 (S.J.R.) F31 MH111108-01A1 (M.F.), T32 MH096678 (M.F.), T32 MH087004 (M.F.), and R01 MH51399 (E.J.N.).

## Author Contributions

Stereotaxic surgeries were performed by M.F., H.A., M.P., and A.T. Immunohistochemistry and *in-situ* hybridization was performed by M.F., C.M., and K.L. Microscopy was performed by M.F., N.T., W.J., and D.B. Molecular cloning of miR-OxR2 constructs was performed by H.A. qPCR was performed by K.C and M.F. Fiber photometry data collection was performed by M.F. and C.J.B and fiber photometry analysis was performed by M.F., C.J.B., E.S.C., and S.B. OxR2 miRNA design was performed by R.J.D. Orexin-cre mouse was made by A.Y. Behavioral experiments were performed by M.F., H.A., L.L, C.J.B, and K.L. Electrophysiology experiments were performed by R.L.C, E.K.L., G.W.H and B.M.A. Results were analyzed and interpreted by M.F. and S.J.R. The manuscript was written by M.F. and S.J.R. and edited by all authors.

---

**Figure S1:**
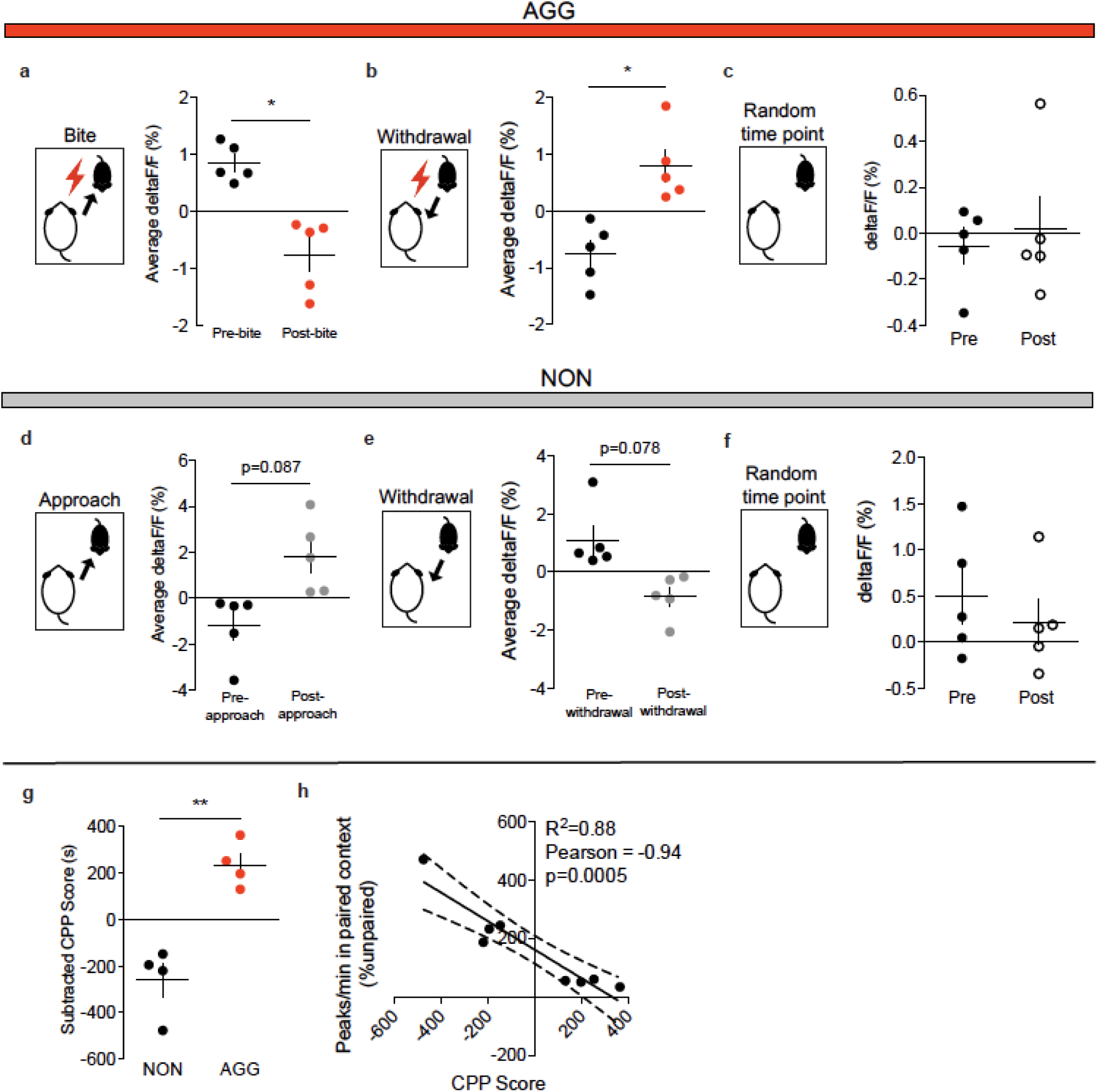
LHb non-conditional fiber photometry supporting data. **a,** AGG average LHb activity was reduced following a bite on day 1 of RI (paired t-test, n=5 mice, 3-5 bites/mouse, t(4)=3.763, p=0.0197). **b,** AGG average LHb activity was increased following a withdrawal from aggression on day 1 of RI (paired t-test, n=5 mice, 3-5 withdrawals/mouse, t(4)=3.229, p = 0.03). **c,** AGG average LHb activity did not differ before and after random time points during RI on day 3 (paired t-test, n=5 mice, t(4)=0.6545, p = 0.5485). **d,** NON average LHb activity was not significantly increased following intruder approach on day 1 of RI (paired t-test, n=5 mice, 3-5 approaches/mouse, t(4)=2.25, p = 0.087). **e,** NON average LHb activity was not significantly reduced following withdrawal from social interactions on day 1 of RI (paired t-test, n=5 mice, 3-5 withdrawals/mouse, t(4)=2.353, p = 0.078). **f,** NON average LHb activity did not differ before and after random time points during RI on day 3 (paired t-test, n=5 mice, 5 time points/mouse, t(4)=0.553, p = 0.6221). **g,** AGGs used for fiber photometry experiments displayed significantly higher aggression CPP scores than NONs (student’s t-test, n=4 mice, t(6)=5.591, p = 0.0014). **h,** LHb peaks in the intruder paired context during the CPP preference test were negatively correlated with CPP score (Pearson correlation coefficient = −0.94, R^2^ = 0.88, p = 0.0005). *p<0.05, **p<0.01. All data are expressed as mean ± SEM.

**Figure S2:**
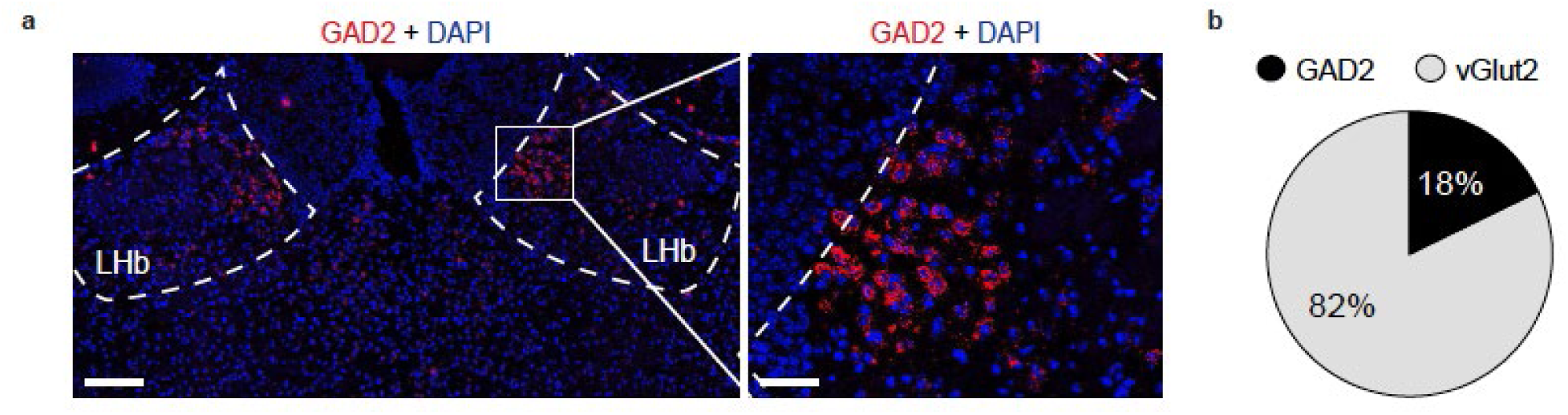
Anatomical characterization of GAD2 LHb neurons. **a,** *in-situ* hybridization (ISH) for GAD2 in the LHb, left image scale bar=150 μm, right image scale bar=70 μm. **b,** Pie chart depicting percentage of LHb neuons that are positive for GAD2 or vGlut2 mRNA as determined by ISH.

**Figure S3:**
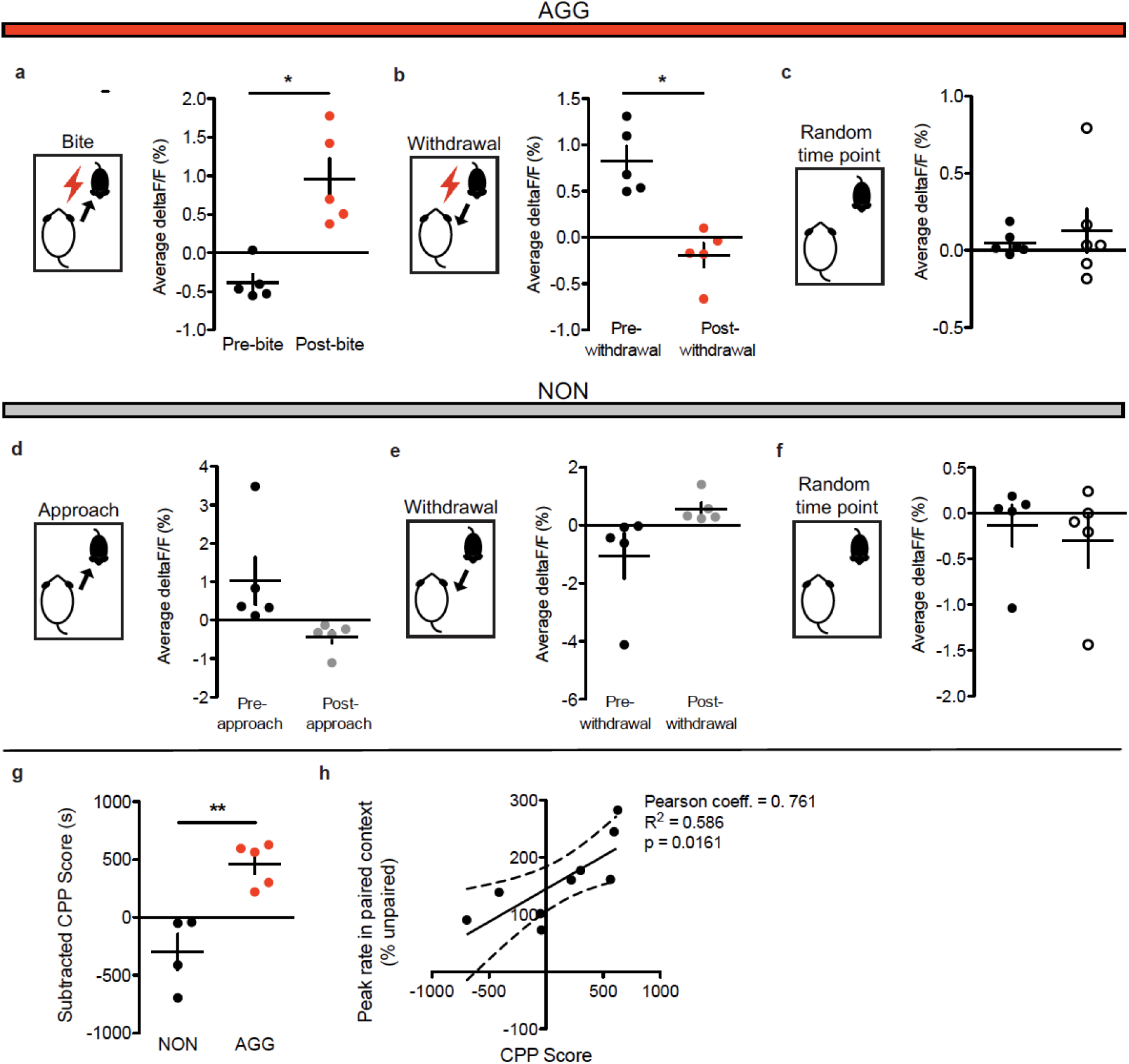
LHb GAD2 neuron fiber photometry supporting data. **a,** AGG LHb GAD2 neuron activity was increased following bites on day 1 of RI (paired t-test, n=5 mice, 2-5 bites per mouse, t(4)=4.008, p=0.016). **b,** AGG LHb GAD2 neuron activity was reduced following a withdrawal from an aggressive encounter on day 1 of RI (paired t-test, n=5 mice, 3-5 withdrawals per mouse, t(4)=3.982, p=0.0164). **c,** AGG LHb GAD2 neuron activity did not differ before and after random times points on day 3 of RI (paired t-test, n=5 mice, 5 time points per mouse, t(4)=0.493, p=0.6475). **d,** NON LHb GAD2 neuron activity was not different before and after an approach on day 1 of RI (paired t-test, n=5 mice, 3-5 approaches per mouse, t(4)=1.843, p=0.1406). **e,** NON LHb GAD2 neuron activity was not different before and after a withdrawal from a non-aggressive social interaction on day 1 of RI (paired t-test, n=5 mice, 3-5 withdrawals per mouse, t(4)-1.633, p=0.1777). **f,** NON LHb GAD2 neuron activity did not differ before and after random time points on day 3 of RI (paired t-test, n=5 mice, t(4)=1.721, p=0.1634). **g,** GAD2-cre AGGs used for fiber photometry experiments displayed significantly higher aggression CPP scores than GAD2-cre NONs (paired t-test, 5=5 mice, t(4)=2.885, p=0.0448). **h,** LHb GAD2 neurons peaks in the intruder paired context during the CPP preference test were positively correlated with CPP score (Pearson correlation coefficient = 0.761m, R^2^=0.586, p=0.0161). *p<0.05, **p<0.01. *p<0.05. All data are expressed as mean ± SEM.

**Figure S4:**
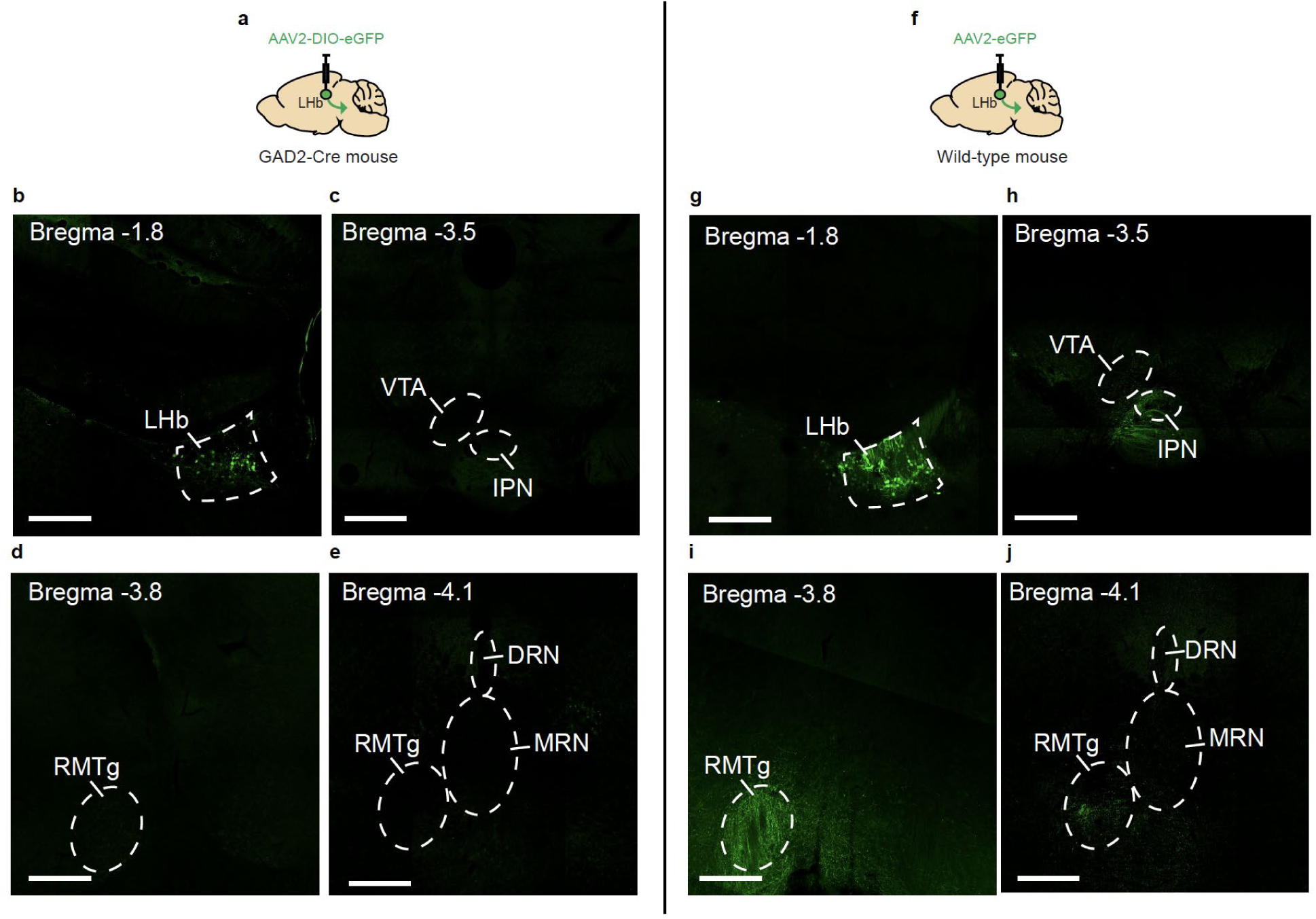
Anterograde tracing of GAD2 LHb neuron projections. **a,** Schematic of surgical manipulations for anterograde tracing of GAD2 LHb neurons. **b,** Representative image of viral infection in GAD2 LHb neurons. **c,** Representative image of the interpeduncular nucleus (IPN) and ventral tegmental area (VTA) in mice expressing eGFP in GAD2 LHb neurons. **d,** Representative image of the rostromedial tegmental nucleus (RMTg) in mice expressing eGFP in GAD2 LHb neurons. **e,** Representative image of the RMTg and anterior dorsal and median raphe nuclei (DRN and MRN) in mice expressing eGFP in GAD2 LHb neurons. **f,** Schematic of surgical manipulations for non-conditional anterograde tracing of LHb neurons. **g,** Representative image of viral infection in LHb neurons. **h,** Representative image of the interpeduncular nucleus (IPN) and ventral tegmental area (VTA) in mice expressing eGFP in LHb neurons. **i,** Representative image of the rostromedial tegmental nucleus (RMTg) in mice expressing eGFP in LHb neurons. **j,** Representative image of the RMTg and anterior dorsal and median raphe nuclei (DRN and MRN) in mice expressing eGFP in LHb neurons. Scale bars= 500 μm.

**Figure S5:**
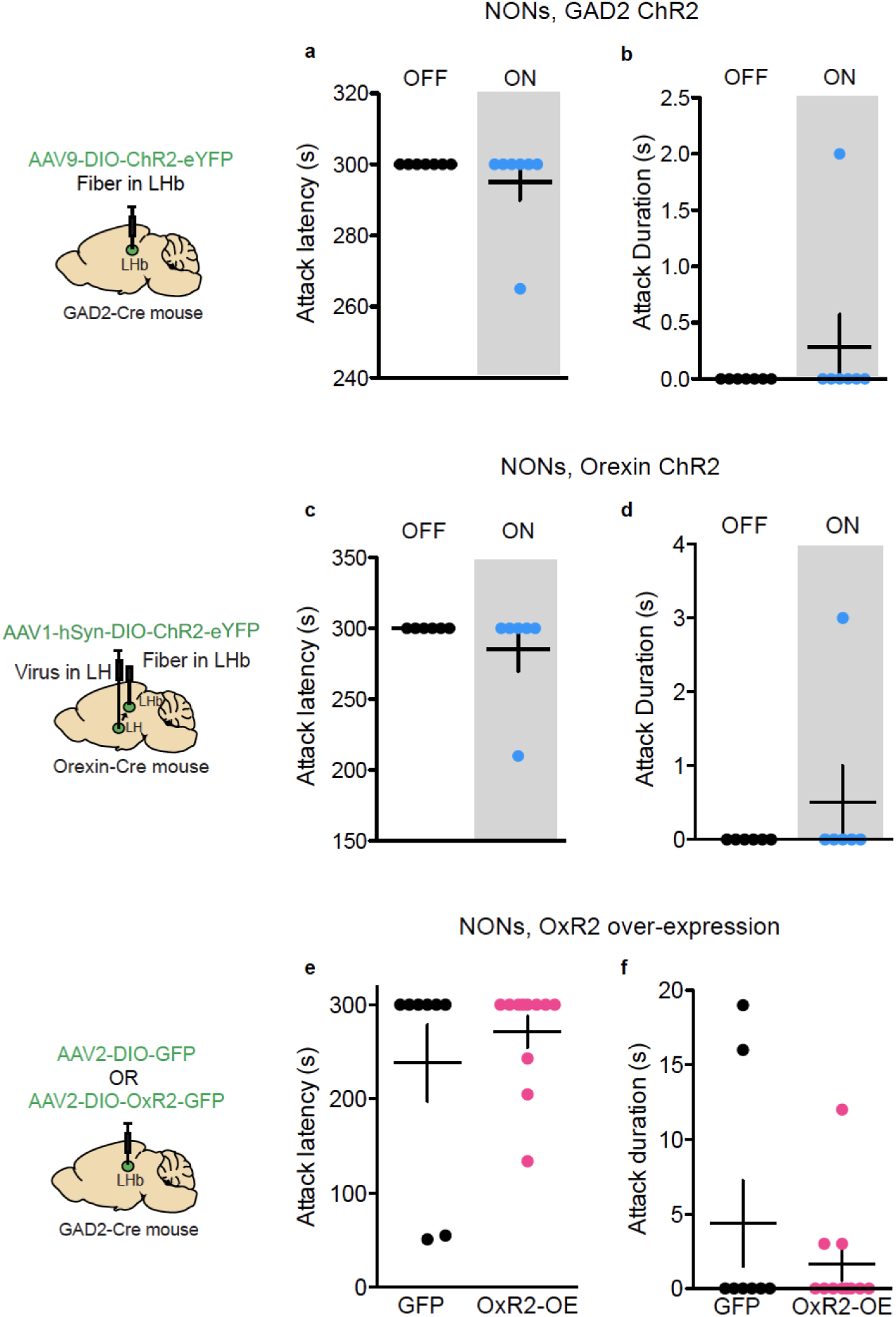
RI behavior in NONs during optogenetic stimulation or OxR2 over-expression in LHb GAD2 neurons. **a,** ChR2-mediated stimulation of GAD2 LHb neurons in NONs did not affect attack latency during RI (paired t-test, n=7 mice, t(6)=1.0, p=0.3559). **b,** ChR2-mediated stimulation of GAD2 LHb neurons in NONs did not affect attack duration during RI (paired t-test, n=7 mice, t(6)=1.0, p=0.3559). **c,** ChR2-mediated stimulation of orexin terminals in the LHb did not affect attack latency during RI (paired t-test, n=6 mice, t(5)=1.0, p=0.3632). **d,** ChR2-mediated stimulation of orexin terminals in the LHb did not affect attack duration during RI (paired t-test, n=5 mice, t(5)=1.0, p=0.3632). **e,** Over-expression of OxR2 in GAD2 LHb neurons in NONs did not affect attack latency during RI (student’s t-test, n=8 GFP and n=11 OxR2-OE, t(17)=0.8338, p=0.4160). **f,** Over-expression of OxR2 in GAD2 LHb neurons in NONs did not affect attack duration during RI (student’s t-test, n=8 GFP and n=11 OxR2-OE, t(17)=0.9951. All data are expressed as mean ± SEM.

**Figure S6:**
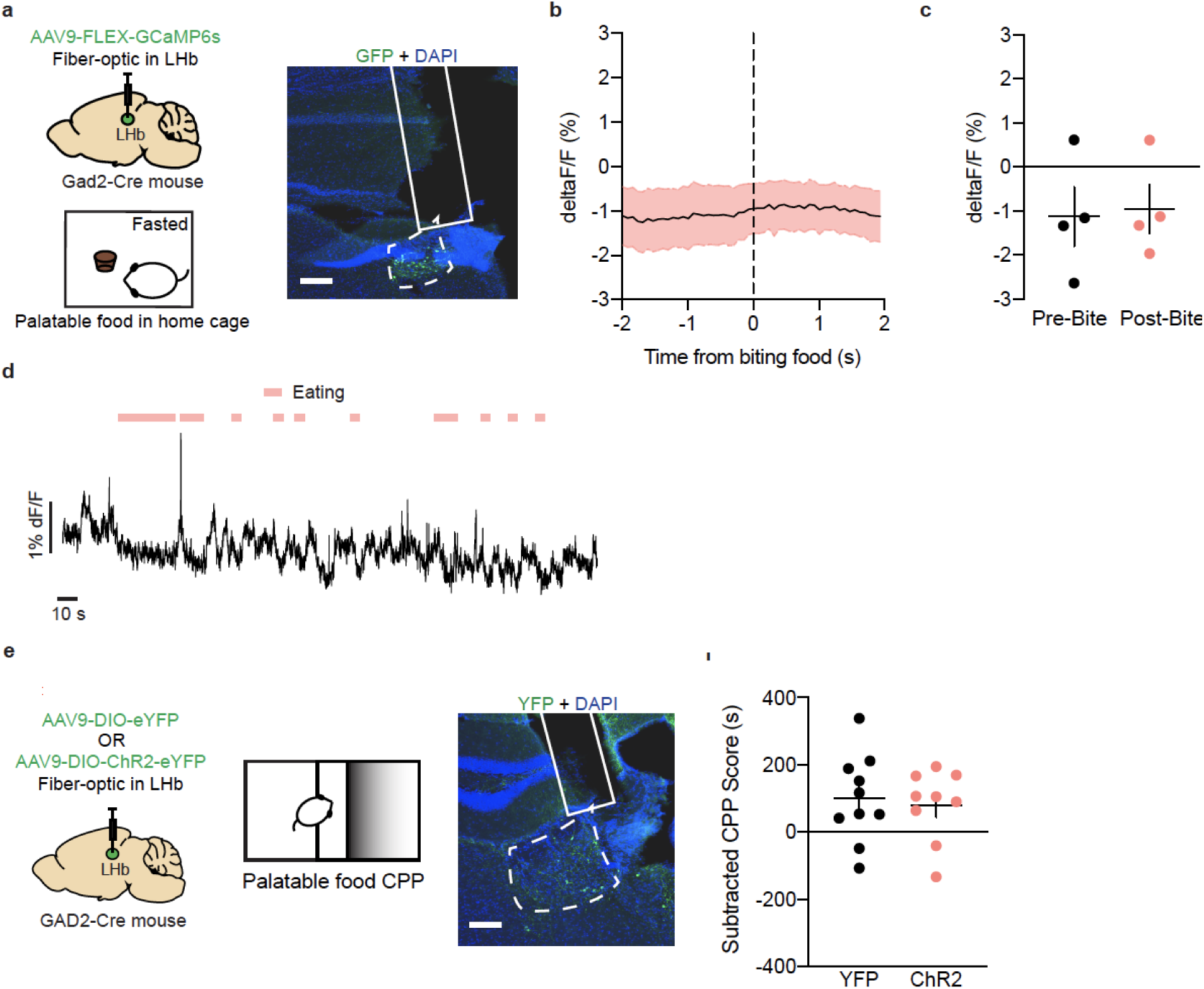
LHb GAD2 neurons are not involved in palatable food reward. **a,** Surgical manipulations and representative viral infection image for LHb GAD2 neuron palatable food reward photometry experiments, scale bar = 400 μm. **b,** Peri-event plot of LHb GAD2 activity 2s before and after biting food in a fasted state in the home cage. **c,** Average LHb GAD2 neuron activity was not different before and after the bite (paired t-test, n=4 mice, t(3)=1.077, p=0.3603). **d,** Representative trace of LHb GAD2 neuron activity during exposure to palatable food in the home cage. **e,** Surgical manipulations and representative viral infection image for LHb GAD2 palatable food CPP optogenetics (ChR2) experiments, scale bar =200 μm. **f,** ChR2-mediated optogenetic stimulation of LHb GAD2 neurons during the palatable food CPP test did not alter the amount of time spent in the palatable food-paired context (student’s t-test, n=10 YFP and n=9 ChR2, t(17)=0.3572, p=0.7253). All data are expressed as mean ± SEM.

**Figure S7:**
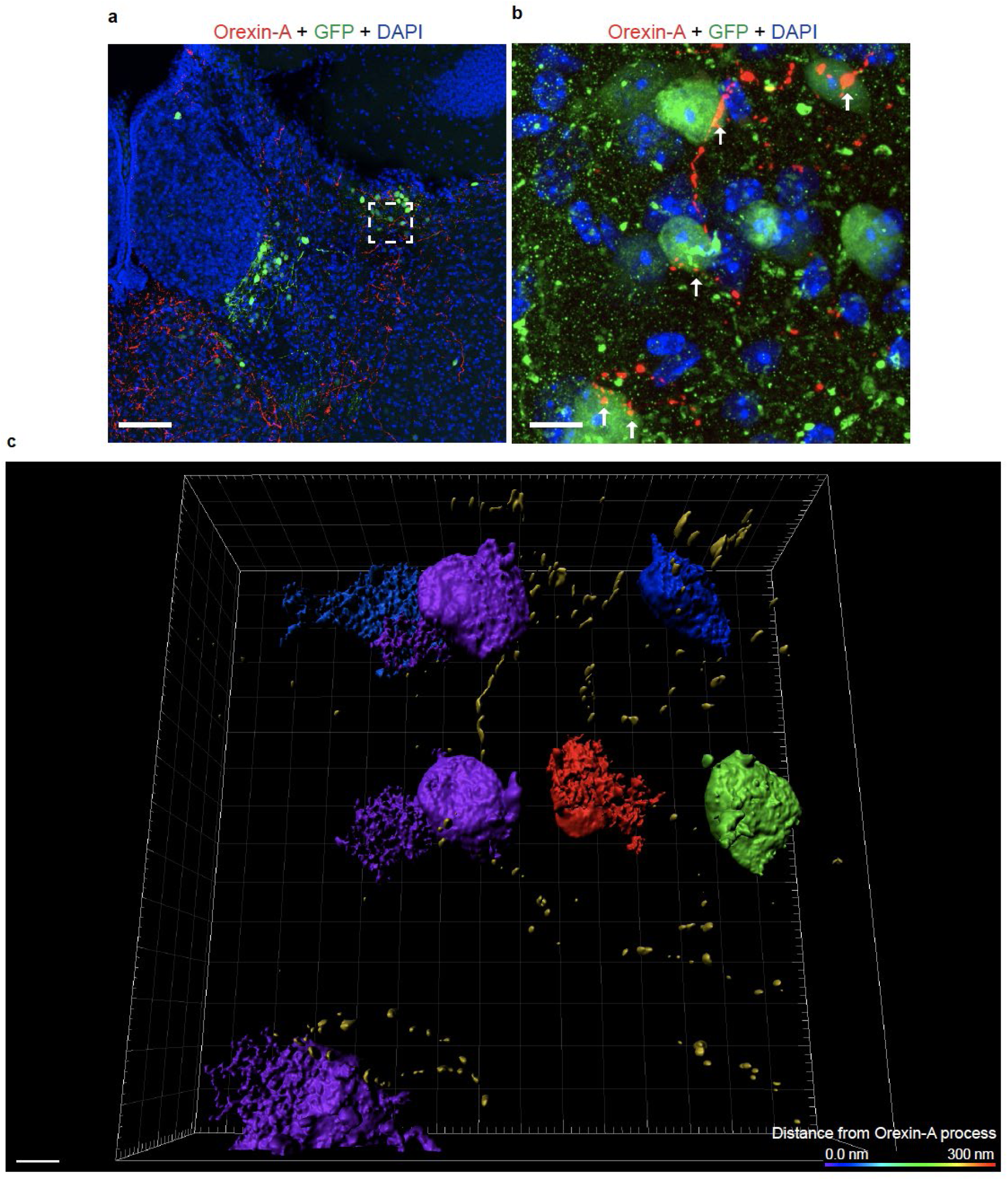
Histology and 3D rendering of GAD2 LHb neurons and orexin axons. **a,** Immunohistochemistry for orexin-A (red), DAPI (blue), and eGFP (green) in a GAD2-Cre mouse injected with AAV-DIO-eGFP, scale bar = 300 μm. **b,** Immunohistochemistry for orexin-A (red), DAPI (blue), and GFP (green) in a GAD2-Cre mouse injected with AAV-DIO-eGFP, scale bar = 10 μm. **c,** 3D rendering of image in b, color of GAD2 neuron coincides with estimated distance from orexin-A axon according to key in lower right corner, scale bar = 5 μm.

**Figure S8:**
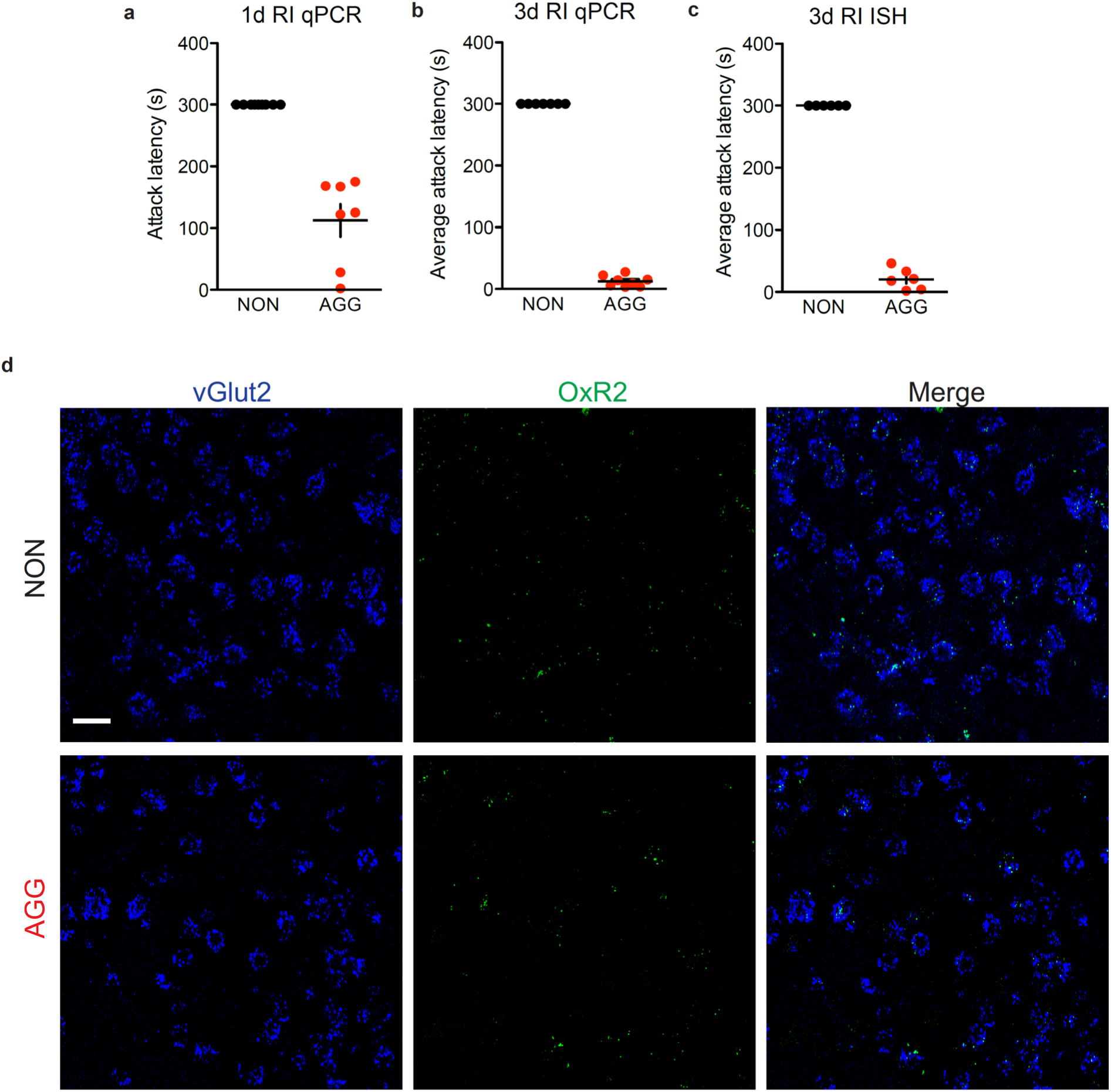
Attack latencies for AGGs and NONs used in qPCR and ISH experiments. **a,** Attack latency for one day of RI in mice used for LHb qPCR. **b,** Average attack latency for three days of RI in mice used for LHb qPCR. **c,** Average attack latency for three days of RI in mice used for LHb OxR2 ISH. **c,** Representative images from OxR2 ISH in AGG and NON LHb vGlut2 neurons following RI, accompanies Fig. 5j, scale bar = 20 μm. Notably, OxR2 expression was barely detectable in these neurons in AGGs or NONs, which is in line with our findings showing low OxR2 expression in vGlut2 neurons in Fig. 5b-c. All data are expressed as mean ± SEM.

**Figure S9:**
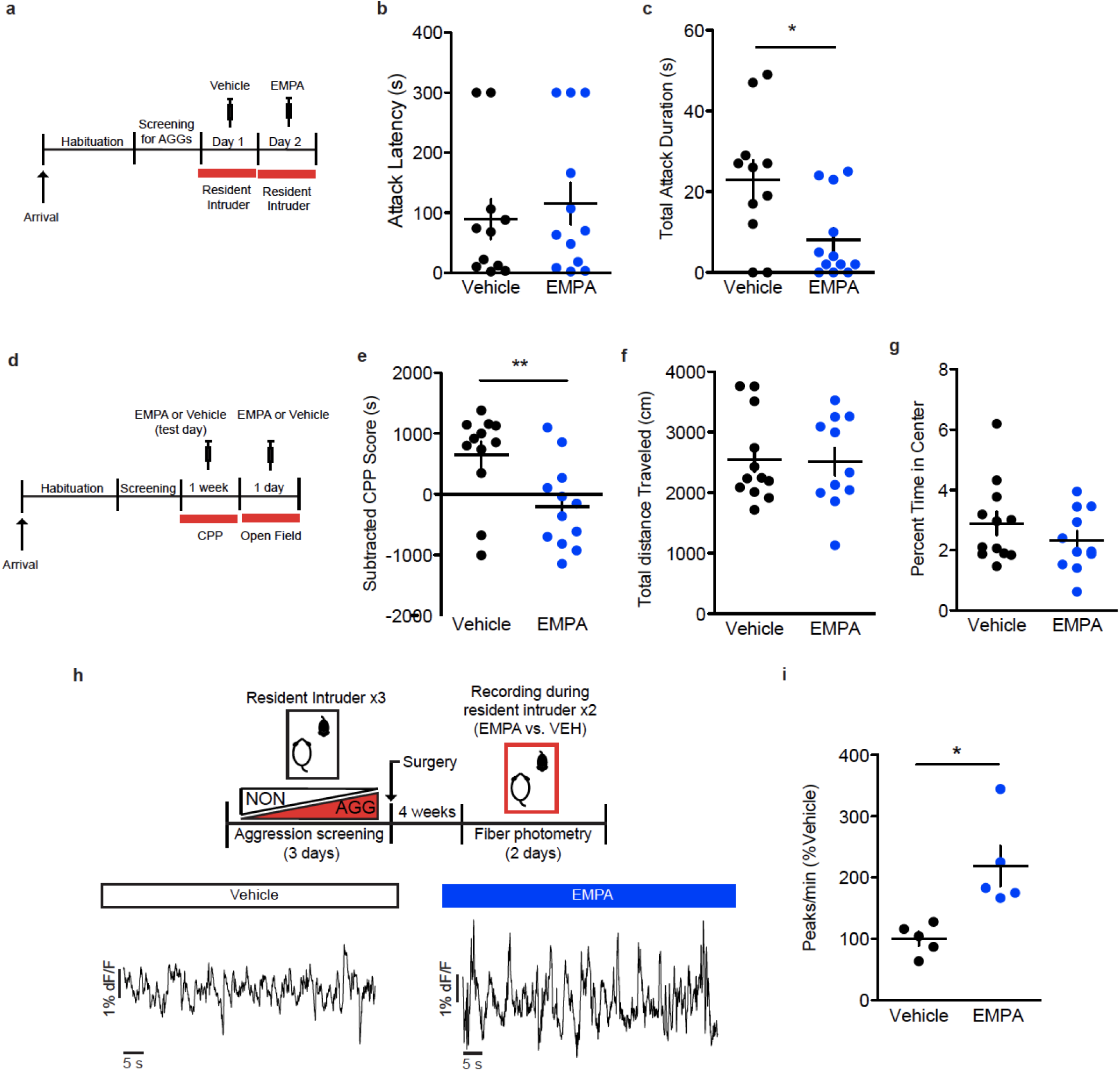
Effects of systemic antagonism of OxR2 with EMPA on aggression. **a,** Experimental scheme for OxR2 systemic antagonism RI experiment. **b,** RI test attack latency in animals treated with EMPA and vehicle (paired t-test, n=11 per group, t(10)=0.3215, p=0.758). **c,** RI test attack duration in animals treated with EMPA and vehicle (paired t-test, n=11 per group, t(10)=2.888, p=0.016).**d,** Experimental scheme for OxR2 systemic antagonism aggression CPP and locomotion experiments. **e,** Aggression CPP for animals treated with EMPA and vehicle (student’s t-test, n=12 vehicle and n=11 EMPA, t(21)=2.885, p=0.0086). **f,** Locomotor activity in the open field for animals treated with EMPA and vehicle (studet’s t-test, n=12 vehicle and n=11 EMPA, t(21)=0.1301, p=0.8991). **g,** Anxiety-related behavior in the open field for animals treated with EMPA or vehicle (student’s t-test, n=11 per group, t(21)=1.134, p=0.2695) **h,** Representative fiber photometry traces in an animal treated with vehicle and EMPA. **i,** LHb GCaMP peaks during RI during vehicle and EMPA treatment (paired t-test, n=5, t(4)=2.946, p=0.0421). *p<0.05, **p<0.01. All data are expressed as mean ± SEM.

**Figure S10:**
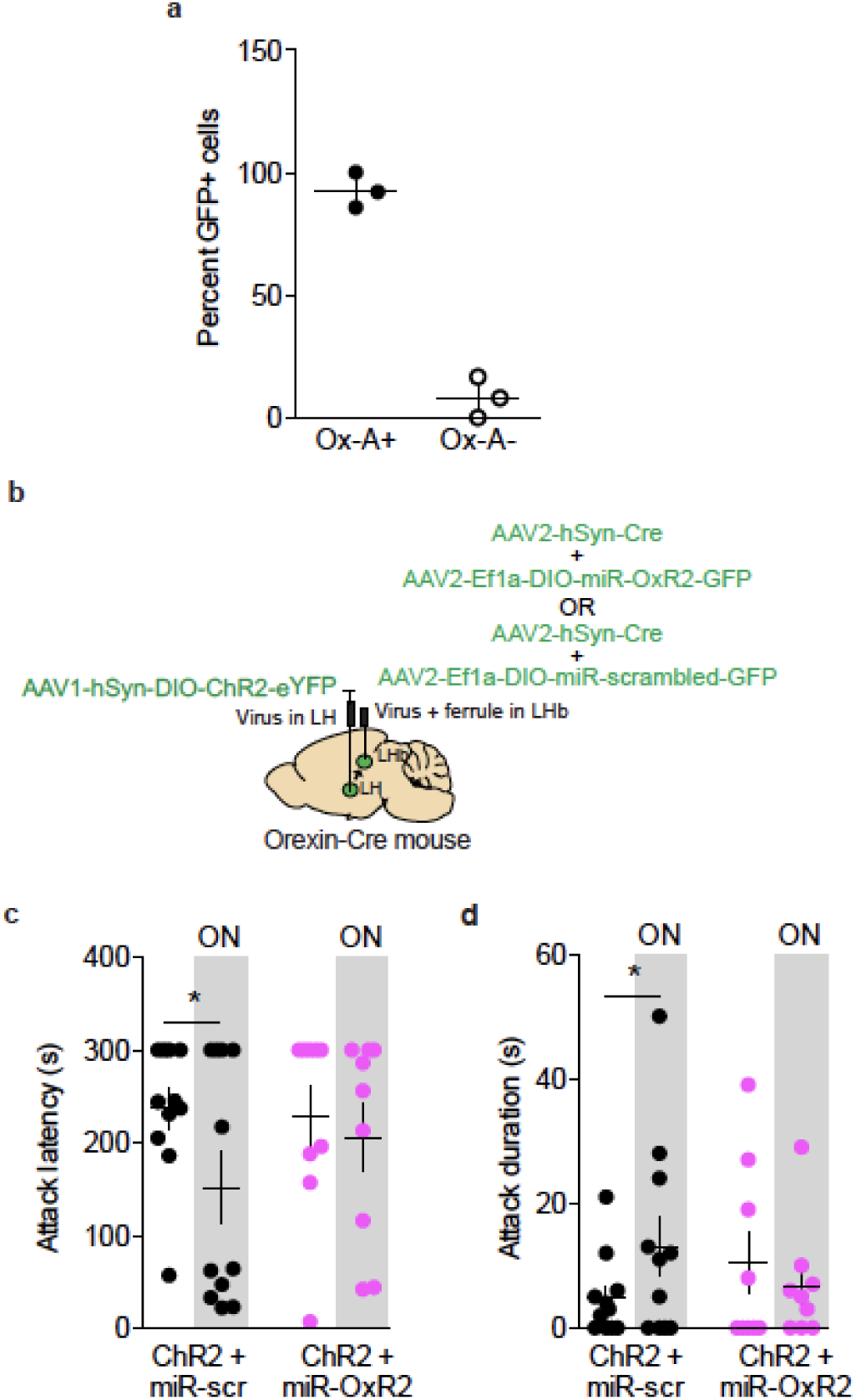
LHb orexin-ChR2 supporting data. **a,** >90% of neurons infected with AAV1-DIO-YFP were positive for orexin-A protein as determined by immunohistochemistry. **b,** Surgical manipulations for ChR2-mediated activation of orexin terminals in the LHb with concurrent knockdown of LHb OxR2. **c,** Optogenetic stimulation of orexin terminals in the LHb reduced attack latency in mice treated with the miR-scrambed virus, but not the miR-OxR2 virus (paired t-test, miR-scrambled: n=11 mice, t(10)=2.424, p=0.0358; miR-OxR2: n=9 mice, t(8)=0.5281, p=0.6117). **d,** Optogenetic stimulation of orexin terminals in the LHb increased attack duration in mice treated with the miR-scrambled virus, but not the miR-OxR2 virus (paired t-test, miR-scrambled: n=11 mice, t(10)=2.260, p=0.0474; miR-OxR2: n=9 mice, t(8)=0.8493, p=0.4204). *p<0.05. All data are expressed as mean ± SEM.

**Figure S11:**
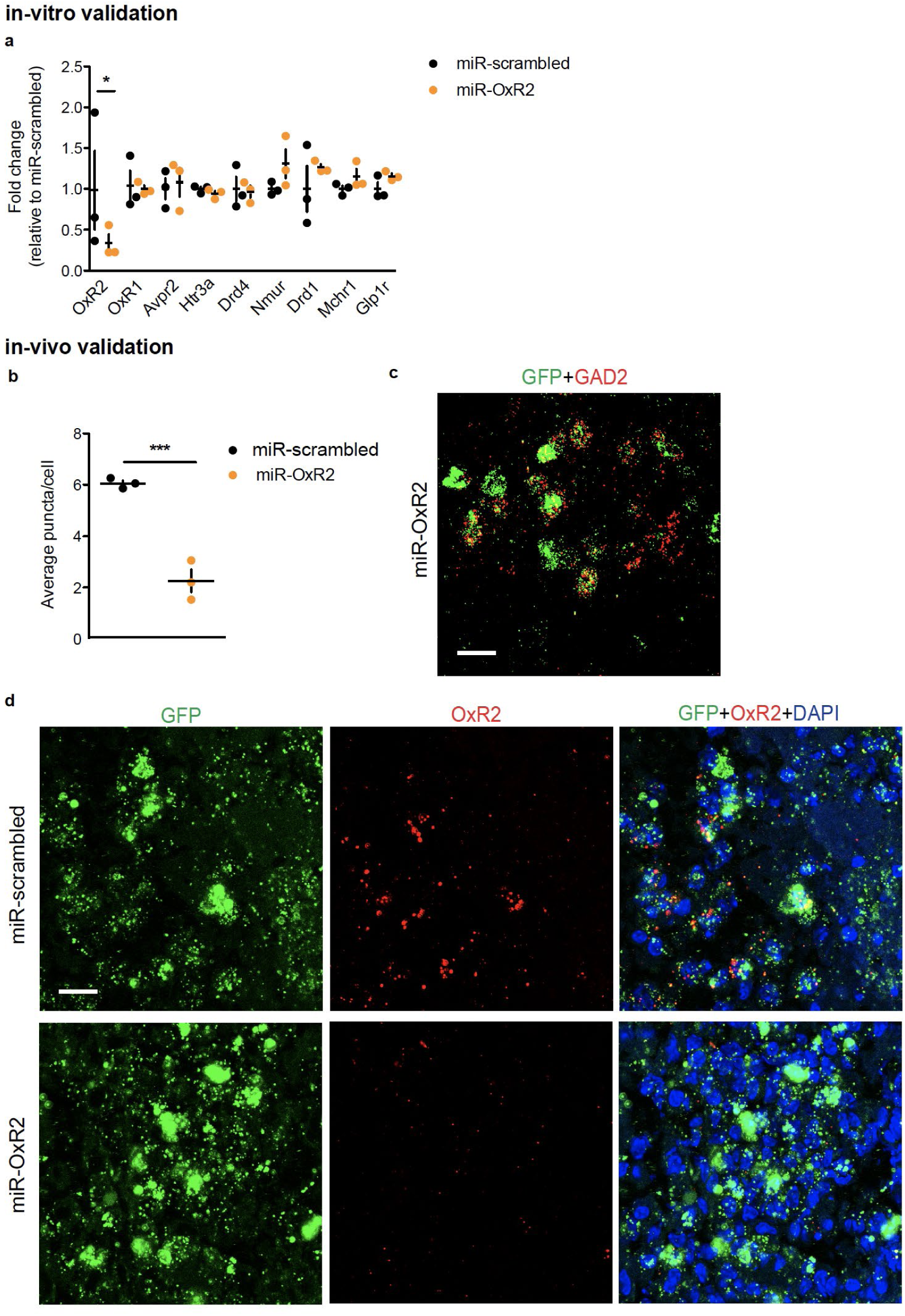
*In-vitro* and *in-vivo* validation of AAV-DIO-miR-OxR2 virus. **a,** N2A cells treated with miR-OxR2 construct selectively reduced OxR2 expression compared to cells treated with miR-scrambled construct, but did not reduce expression of related transcripts (student’s t-test, n=3 per group; OxR2: t(4)=2.402, p=0.0482; OxR1: t(4)=0.2123, p=0.8423; Avpr2: t(4)=0.3686, p=0.7311; Htr3a: t(4)=1.309, p=0.2607; Drd4: t(4)=0.1925, p=0.8567; Nmur: t(4)=1.672, p=0.1699; Drd1: t(4)=0.9239, p=0.4078; Mchr1: t(4)=1.467, p=0.2163; Glpr1: t(4)=1.785, p=0.1488). **b,** GAD2-Cre mice injected with AAV-DIO-miR-OxR2 displayed reduced expression of OxR2 compared to mice injected with AAV-DIO-miR-scrambled as determined by ISH (student’s t-test, n=3 mice, 2 slices per mouse, t(4)=18.44, p=0.0001). **c,** Representative image of GFP expression localized to GAD2 LHb neurons in GAD2-Cre mice injected with AAV-DIO-miR-OxR2, scale bar = 25 μm. **d,** Representative images of OxR2 expression in GAD2 LHb neurons infected with AAV-DIO-miR-OxR2 or AAV-DIO-miR-scrambled, scale bar, 20 μm. *p<0.05, ***p<0.001. All data are expressed as mean ± SEM.

**Figure S12:**
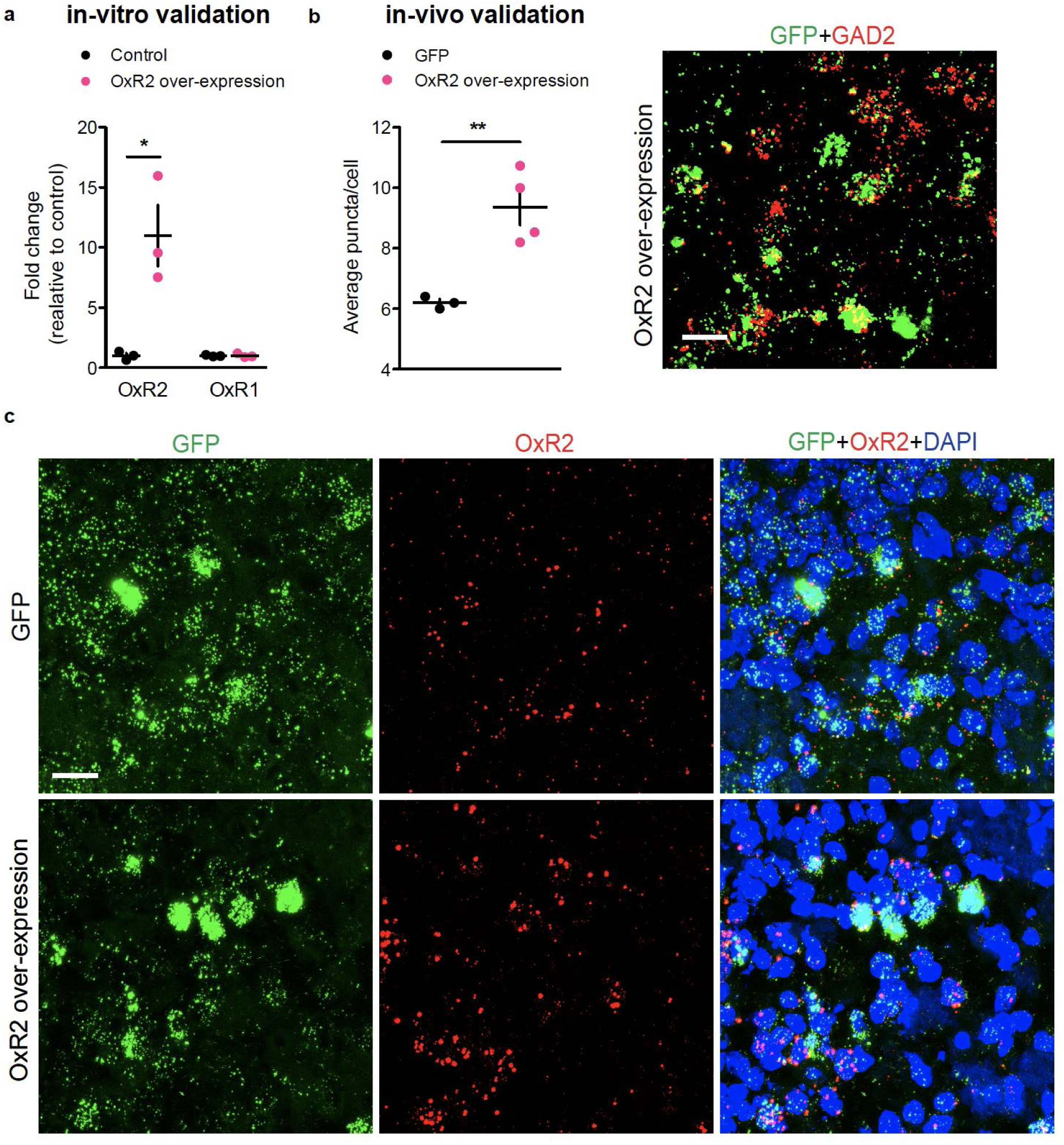
*In-vitro* and *in-vivo* validation of AAV-DIO-OxR2 virus. **a,** N2A cells treated with OxR2 over-expression construct selectively increased OxR2 expression compared to controls (student’s t-test, n=3 per group, OxR2: t(4)=3.939, p=0.0171; OxR1: t(4)=0.1238, p=0.9075). **b,** GAD2-Cre mice injected with AAV-DIO-OxR2 displayed increased expression of OxR2 compared to mice injected with AAV-DIO-GFP as determined by ISH (student’s t-test, n=3 mice, 2 slices per mouse, t(4)=4.417, p=0.0069) (left). Representative image of GFP expression localized to GAD2 LHb neurons in GAD2-Cre mice injected with AAV-DIO-OxR2, scale bar = 25 μm (right). **c,** Representative images of OxR2 expression in GAD2 LHb neurons infected with AAV-DIO-OxR2 or AAV-DIO-GFP, scale bar = 25 μm. *p<0.05, **p<0.01. All data are expressed as mean ± SEM.

**Figure S13:**
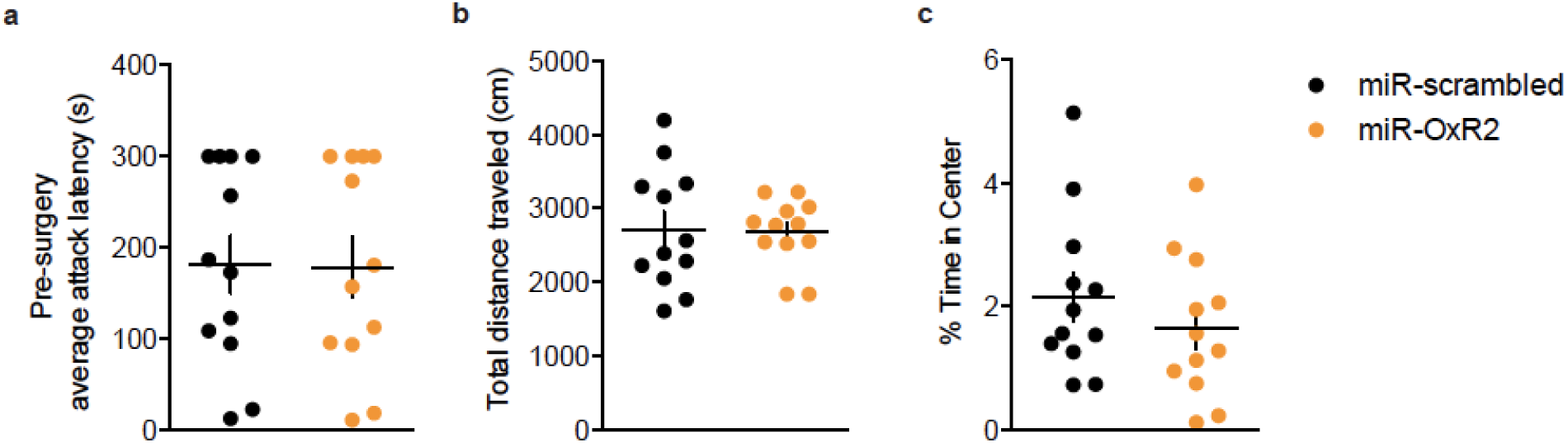
Pre-surgery attack latency, locomotor behavior, and anxiety-related behavior for GAD2-cre mice treated with miR-scrambled or miR-OxR2 viruses. **a,** miR-scrambled and miR-OxR2 mice did not display differences in attack latency before surgery (student’s t-test, n=12 per group, t(22)=0.0693, p=0.946). **b,** Following surgery, miR-OxR2 mice did not display differences in total distance traveled in the open field compared to miR-scrambled mice (student’s t-test, n=12 per group, t(22)=0.1639, p=0.8713). **c,** miR-OxR2 mice did not display any differences in anxiety-related behavior in the open field compared to miR-scrambled mice (student’s t-test, t(22)=1.012, p=0.3224. All data are expressed as mean ± SEM.

